# Acetylation regulates the oligomerization state and activity of RNase J, the major ribonuclease of *Helicobacter pylori*

**DOI:** 10.1101/2021.08.09.455701

**Authors:** Alejandro Tejada-Arranz, Maxime Bouilloux-Lafont, Xue-Yuan Pei, Thibaut Douché, Mariette Matondo, Allison Williams, Bertrand Raynal, Ben F Luisi, Hilde De Reuse

## Abstract

In *Helicobacter pylori*, post-transcriptional regulation strongly relies on the activity of an RNA degradosome, composed of the essential ribonuclease RNase J and the DEAD-box RNA helicase RhpA. Here, we describe post-translational modifications of this protein complex that affect its activity. Cell-extracted RNase J is acetylated on multiple residues, one of which, K649, strongly impacts RNase J oligomerization, which in turn influences ribonuclease activity. Corroborating the link between oligomerization and activity, mutations targeting K649 and other residues affect the dimerization and *in vitro* activity of RNase J. Our crystal structure of RNase J reveals three loops that gate access to the active site and rationalizes how oligomerization state influences activity. The acetylated residues of RNase J are important for *H. pylori* morphology, highlighting that the modifications affect the RNase J cellular function. We propose acetylation as a regulatory level controlling the activity of RNase J and the *H. pylori* RNA degradosome.

## Introduction

Post-transcriptional regulation is one of the most important levels of control of gene expression in every kingdom of life. Ribonucleases (RNases) are key enzymes in this regulation, that are involved in RNA maturation and degradation. In several bacteria, major RNases act in concert with DEAD-box RNA helicases, that help to unfold RNA secondary structures, within protein machines called RNA degradosomes^1^.

We previously demonstrated the existence of a minimal RNA degradosome in *Helicobacter pylori*^2^. This Gram-negative bacterial pathogen chronically colonizes the stomach of half of the human population worldwide. Infection leads to chronic gastritis and, in some cases, to further gastric pathologies such as peptic ulcers, MALT lymphoma, or gastric adenocarcinoma that causes 800,000 deaths each year worldwide^3^. *H. pylori* possesses a small genome (1.6 Mb) with few transcriptional regulators, and several lines of evidence indicate that post-transcriptional regulation plays a major role in gene expression control in this organism^4–6^. The *H. pylori* RNA degradosome is composed of RNase J, a 77.6 kDa enzyme that possesses both 5′-3′ exoribonuclease and endoribonuclease activities, and RhpA, a DEAD-box RNA helicase, with a molecular weight of 55.8 kDa^2, 7^. RNase J is essential in *H. pylori* and plays a major role in the degradation of both mRNAs and antisense RNAs ^8^. RhpA is the sole DEAD-box RNA helicase encoded by the *H. pylori* genome and is essential for colonization in the mouse infection model^7^. Furthermore, we recently demonstrated that, in *H. pylori*, the RNA degradosome is compartmentalized into clusters at the inner membrane whose formation is regulated and likely represents a major level of control of its activity^9^.

RNase J-encoding genes are present in 57 % of sequenced bacteria^1^, including the Gram-positive organisms *Bacillus subtilis*, *Staphylococcus aureus* and *Deinococcus radiodurans*. *B. subtilis* and *S. aureus* encode two paralogues of RNase J, called J1 and J2 while *H. pylori, Streptomyces pyogenes* and *D. radiodurans* possess only one gene for this protein. Compared to the RNase J proteins of these three organisms, the *H. pylori* enzyme possesses a unique N-terminal extension of 132 residues that is predicted to be predominantly disordered^2, 9^. In *B. subtilis*, while both RNase J paralogues (*Bsu*RNases J1 and J2) have equivalent endoribonuclease activities, *Bsu*RNase J2 seems to lack the 5’-3’ exoribonuclease activity that predominates in *Bsu*RNase J1^10^. Furthermore, these two proteins interact and form a heterotetramer *in vitro* that has different cleavage specificity *in vitro* than each protein alone^10^. *In vitro* analysis of the RNase J1 enzyme from *S. aureus* showed that it predominantly assembles into dimers, but also forms tetramers, independently of divalent cations^11^. The single RNase J from *S. pyogenes* forms dimers and tetramers *in vitro* without divalent cations being observed in the dimerization surface^12^. In *D. radiodurans*, RNase J forms a dimer in a Mn-dependent fashion *in vitro*, and the interaction between the two monomers is mediated by the C-terminal domain of *Dra*RNase J^13^. Several crystal structures of RNase J proteins have been published^12–14^, where RNase J is generally found to be a tetramer (dimer of dimers). However, it is unknown whether the different oligomeric states of RNase J detected *in vitro* have differences in activity.

Almost nothing is known about post-translational modifications of these protein complexes and whether this might influence their activity. Acetylation has emerged as a major post-translational regulatory mechanism in bacteria^15, 16^. Although the acetylation of lysine, cysteine, serine and threonine residues has been reported, lysine acetylation is the most prominent modification, particularly on its ε-amino group^15^. This modification has been shown to regulate multiple properties of the target proteins, including enzymatic activity, DNA-binding capability or stability, among others^16^. Interestingly, some proteins involved in RNA metabolism have been reported to be lysine acetylated, including (i) *Escherichia coli* RNase R, whose acetylation regulates its stability^17^; (ii) *E. coli* RNase II, regulated at the activity level^18^; and (iii) the CshA DEAD-box RNA helicase from *B. subtilis*, where acetylation has been proposed to influence its interaction with RNA polymerase independently from its association with RNase Y, the major ribonuclease of the *B. subtilis* RNA degradosome^19^.

The central importance of RNase J in post-transcriptional regulation in *H. pylori* and its participation in an RNase J-based degradosome prompted us to investigate the structure, oligomeric state and post-translational regulation of the *H. pylori* RNase J and to assess its stoichiometry in the RNA degradosome. Here, we report the crystal structure of *H. pylori* RNase J without its N-terminal extension. We also show that RNase J is in equilibrium between monomeric and dimeric states *in vitro* and that, together with its monomeric RNA helicase partner RhpA, assembles into a minimal degradosome complex with a 1:1 stoichiometry. Our data suggest that the dimeric form constitutes the active form of RNase J *in vitro*. Furthermore, we found for the first time that RNase J is acetylated at several residues by multiple mechanisms. We concluded that the acetylation state regulates the oligomerization state and activity of RNase J *in vitro* and its activity in *H. pylori*, and thereby constitutes an important level of control of the RNA degradosome.

## Results

### Structure of the *H. pylori* RNase J

To better understand the structure-function relationship of the *H. pylori* RNase J protein, we first decided to determine its crystal structure. Several attempts to purify sufficient amounts of full-length RNase J from *H. pylori* to obtain crystals were unsuccessful. As we previously reported^9^, the N-terminal part of RNase J (first 120 amino acids) that is specific of the *Helicobacter* species is predicted to be an intrinsically disordered region (IDR). After truncating this region, well diffracting crystals were obtained from this RNase J form comprising residues 137-691 (from *H. pylori* strain 26695, uniprot P56185). The crystal structure of *H. pylori* RNase J was solved at 2.75 Å by molecular replacement using the *Streptomyces coelicolor* homologue as a search model (Table S1). One molecule occupies the asymmetric unit of the crystal, and through crystal symmetry a homotetramer can be generated that is organized as a dimer-of-dimers (Figure 1A). The core region of the enzyme, residues 137-586, encompasses the beta-CASP and beta-lactamase domains that define this family^20^, but there are no detectable metal cofactors in the map at the catalytic site. An X-ray Zn-peak scan also confirmed that there are no detectable Zn ions in the crystal.

**Figure 1.**
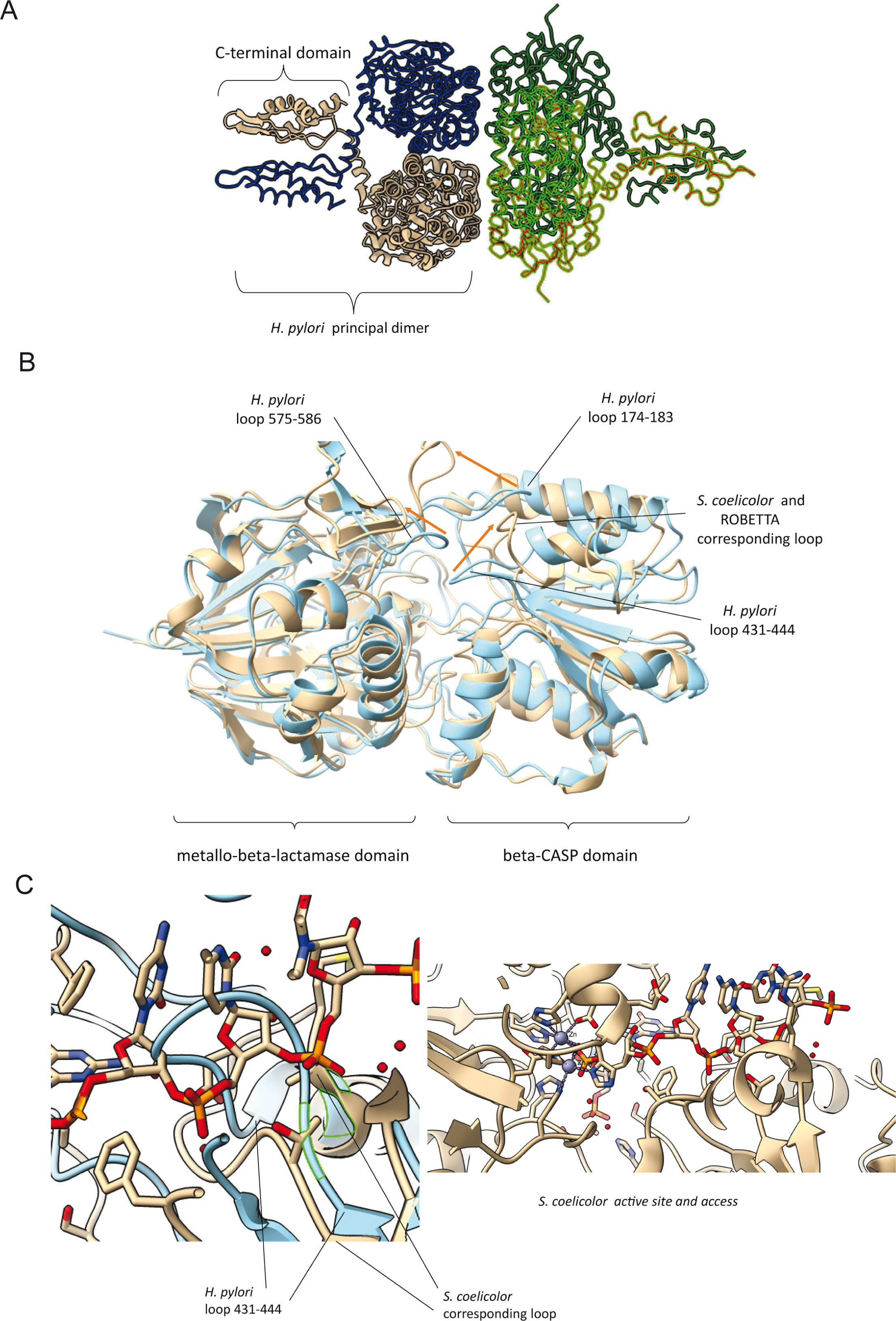
Structure of RNase J from *H. pylori.* Crystals were prepared from *H. pylori* RNase J residues 139-691, encompassing the beta-CASP, metallo-beta-lactamase and C-terminal domains. (A) View of the tetramer, which is a dimer-of-dimers. A monomer occupies the asymmetric unit, and the tetramer is generated through crystallographic symmetry. (B) Overlay of the predicted structure from Robetta server (beige) and the refined crystal structure (blue). The individual beta-CASP and metallo-beta-lactamase domains overlay well, but there are noted differences in the orientation of the three loop regions 174-183, 431-441 and 575-586 at the entrance to the active site (shifts indicated by orange arrows). The predicted structure is closer to the orientation seen in *Streptomyces coelicolor* RNase J, shown in overlay with *H. pylori* enzyme in the left panel of (C). The *S. coelicolor* enzyme structure (beige) is in complex with RNA, and the path of the RNA substrate along a channel to the active site is shown in the right panel. The *H. pylori* RNase J (blue) cannot bind RNA with the same orientation seen for the *S. coelicolor* enzyme due to steric clashes with the loops (left panel). The two zinc ions at the active site of the *S. coelicolor* enzyme are indicated in the right panel.

**Table 1.**
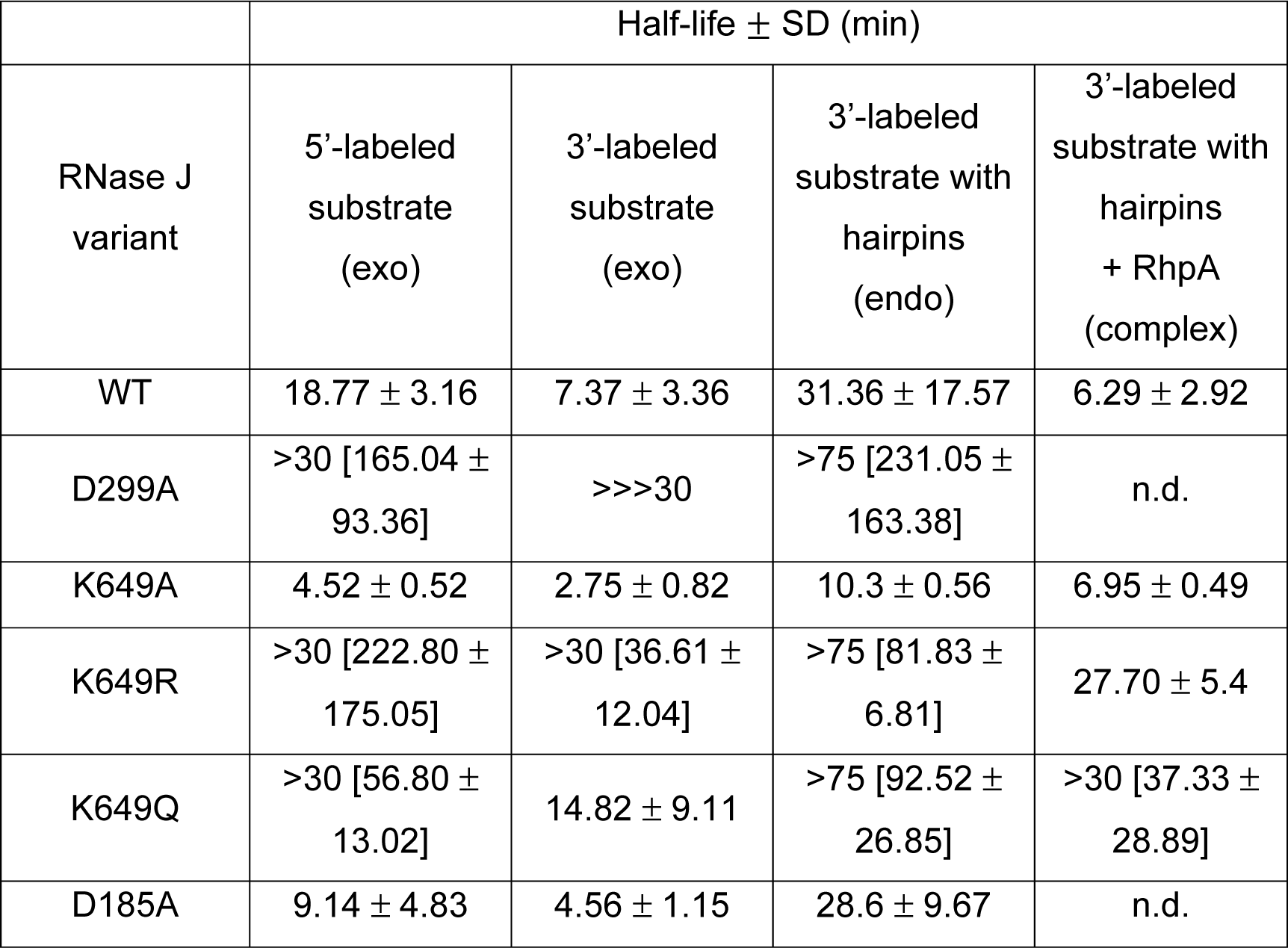
Half-lives of fluorescent RNA substrates in the presence of different variants of RNase J alone or in complex with RhpA. The half-live values in brackets are estimated values extrapolated from the decay slope when the half-lives exceeded the time of the assay, over 30 min for the two first and the last columns and over 75 min for the third column.

The structure predicted by the Robetta server, based on machine learning^21^, is in excellent agreement with the X-ray model for the core, with the notable exception of the three loop regions 174-183, 431-441 and 575-586 at the entrance to the active site, for which the predicted loop positions are closer to those seen in the crystal structure of the *S. coelicolor* RNase J (Figure 1B-C).

The electron density over these regions is well defined in the *H. pylori* map and modelled with high confidence. Comparing the structures, it is apparent that the position of the loops will enforce a different path for the RNA substrate in *H. pylori* RNase J compared with the *S. coelicolor* enzyme (Figure 1C).

The C-terminal domain (CDT), from residues 587 to 691, is poorly ordered in the crystal lattice and the density was not sufficiently well resolved to model the polypeptide. The Robetta model for the C-terminal domain fits well and fitted into the map, showing good agreement with the predicted helical segments. The predicted model is also in agreement with a high confidence homology model prepared by PHYRE2 using the *Thermus thermophilus* homolog as template (PDB 3BK2). This domain makes a self-complementary interaction through crystal lattice contact. The Robetta model predicted with low confidence an extended helical region of the N-terminal residues 1-136 that were not included in the crystallized protein, and this region seems unlikely to have a compact fold. The final model to fit the X-ray data is a fusion of the fitted crystal structure of the core and the Robetta model for the C-terminal domain with side chains removed due to poor resolution of this part of the electron density map.

In the dimeric unit, the protomer interfaces are formed by self-complementary interactions made mostly by the CTD. The surface area buried at the CTD-CTD interface is estimated to be 1165 Å^2^. The beta-CASP and beta-lactamase domains of RNase J also make a self-complementary interface in the principal dimer, with an estimated area of 672 Å^2^ that is a little more than half of the CTD-CTD contribution. Stronger interactions typically have a surface area of more than 2000 Å^2 22^. The interface between the beta-CASP and beta-lactamase domains is very similar to that seen in other dimers of RNase J, for instance in the homologue of *S. coelicolor*^12^. The interfaces that are formed between the dimer units to make the tetramer are also made by the beta-CASP and beta-lactamase interfaces, and these have a total buried area of 2070 Å^2^.

An overlay of RNase J of *H. pylori* and of *S. coelicolor*, where an RNA substrate is bound, suggests that the path of the RNA to the active site cannot be the same for the two enzymes, and that the nucleic acid would need to be rerouted in the *H. pylori* enzyme. The obstructions are caused by exposed peptide loop regions that are positioned to accommodate the CTD. It is possible that changes in the oligomerization state will impact on the loops and access to the active site.

It is notable that the CTD has extensive charged side chains on the surface of proteins that likely form complementary charge interactions. Such an organization may stabilize the fold of the small domain^23–25^. As the self-complementary interactions of the C-terminal domain are a key component of the assembly, any changes in the stability of the CTD may impact on the oligomeric state. This prompted us to investigate the *in vitro* oligomeric state of RNase J and its RhpA partner.

### *In vitro* analysis of the oligomeric state of the two partners of the *H. pylori* RNA degradosome

The crystal structure of *H. pylori* RNase J and of several other RNase J proteins correspond to tetramers. However, for some of these proteins, other oligomeric states (dimers and monomers) were found in solution and the activity of the different forms was unclear. To evaluate the oligomeric state of *H. pylori* RNase J and RhpA *in vitro*, we performed analytical ultracentrifugation (AUC) with His-tagged versions of RNase J and RhpA purified from *E. coli* (Supp Figure S1). The sedimentation coefficient distribution of RNase J indicates that this protein forms two oligomeric species *in vitro* with concentration-independent sedimentation coefficients (S_0_) of 5.8 and 8.8 (Fig. 2A). Based on the structural data of *H. pylori* RNase J presented above, the calculation of the sedimentation coefficients with the HYDROPRO software was 5.4 and 8.9 for a monomer and dimer, respectively. Moreover, the calculated sedimentation for the monomer suggests that the C-terminal end of the monomer is probably flexible and less extended in solution, explaining the small difference in sedimentation coefficient. On the other hand, the calculated sedimentation for the dimer is fully compatible with the extended arrangement of the C-terminal end, showing that dimerization locks the structure in an extended conformation. Therefore, we concluded that the two forms of RNase J observed by AUC are indeed a monomer and a dimer (Figure 2A) that have a structural arrangement in solution which is compatible with the observed crystal structure. The monomeric population predominates (∼60%) over the dimer (∼40%) independently of the protein concentration, showing that these species were not in equilibrium but were dependent on other modifications. AUC of RhpA shows behavior as a monomer independent of concentration with an S_0_ of 3.85 (Figure 2B).

**Figure 2.**
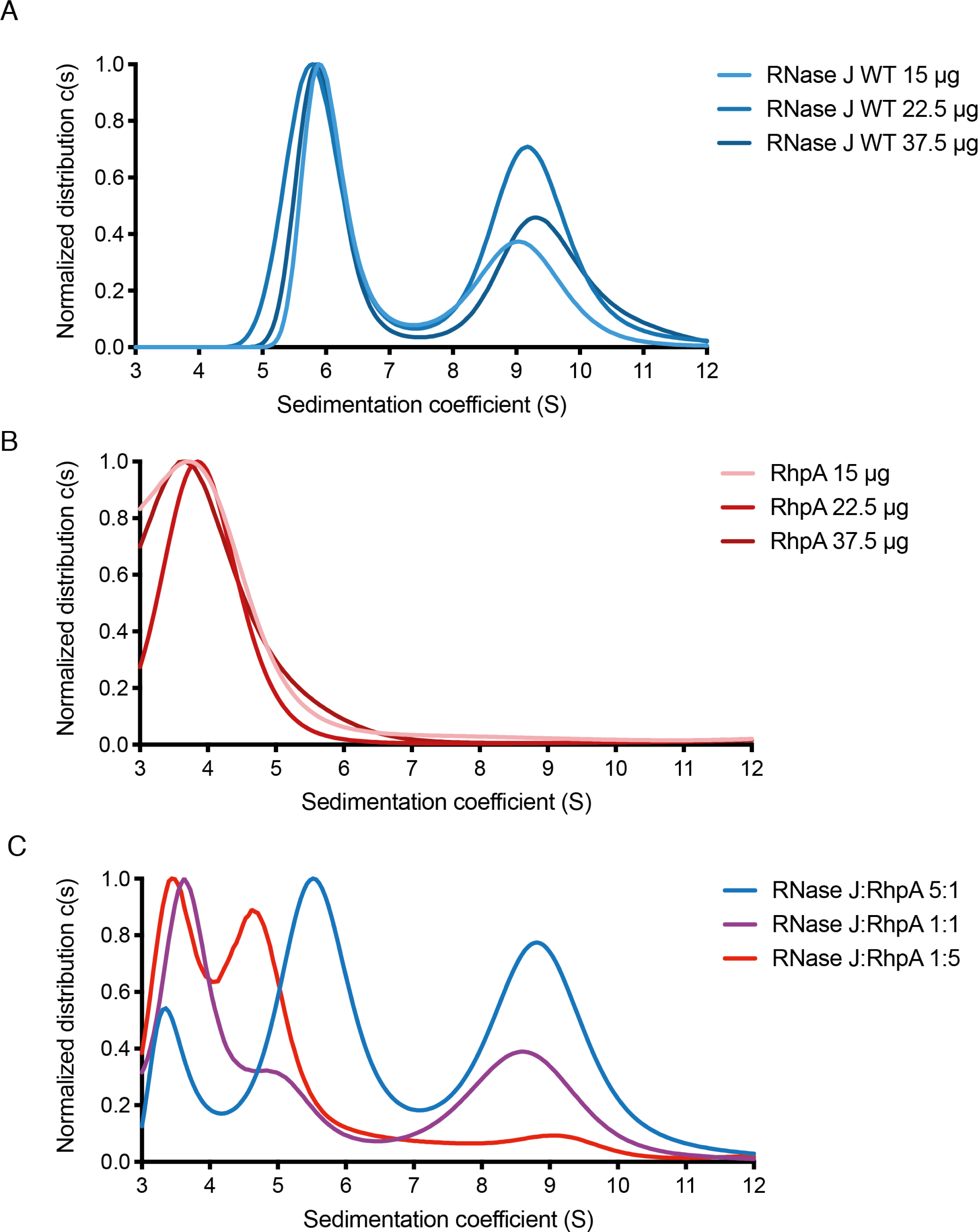
The solution oligomerization state of RNase J, RhpA and the degradosome assembly. The profiles are from analytical ultracentrifugation sedimentation of (A) RNase J, (B) RhpA and (C) the degradosome complex at different ratios of RNase J and RhpA. The profiles were recorded at 230 nm wavelength.

### RNase J and RhpA form a 1:1 minimal degradosome complex

Next, we wanted to evaluate the stoichiometry of the protein complex formed by RNase J and RhpA. We mixed both proteins at 5:1, 1:1 and 1:5 ratios and subjected them to AUC (Figure 2C). When using a 5:1 RNase J:RhpA ratio, we identified three peaks corresponding to both proteins alone (RNase J monomer and dimer and RhpA monomer). With a 1:1 RNase J:RhpA ratio, three peaks could be detected. Based on the previous calculations, these peaks are proposed to correspond to (i) RhpA monomer alone for the 3.85S peak, (ii) a dimer of RNase J for the 8.8S peak, and (iii) a minor peak with a sedimentation coefficient of 4.8S that would correspond to the RNase J-RhpA complex. This latter peak was much more prominent when using a 1:5 ratio of RNase J:RhpA, where only two peaks could be detected, corresponding to RhpA alone and most probably the complex with a coefficient of 4.8S. From this peak and the available frictional coefficients, we calculated a mass for the complex of approximately 124 kDa that is only compatible with a complex with a 1:1 stoichiometry. Interestingly, neither the monomer nor the dimer of RNase J was detected under this condition, and all RNase J was involved in complex formation. We thus conclude that RNase J and RhpA form a 1:1 stoichiometry complex *in vitro*.

### Bacterial two hybrid (BACTH) to identify residues important for the oligomerization of RNase J

As AUC results show that the formation of oligomer is not an equilibrium and is independent of concentration, we then searched for the potential residues involved in the self-interaction of RNase J using bacterial two hybrid (BACTH) experiments that test the interaction of protein pairs in *E. coli*. First, a consistent interaction of wild type RNase J with itself was detected (Figure 3A). Then, different RNase J mutants were constructed and tested. As previously reported for other RNase J proteins^13^, we found that the C-terminal region of RNase J is essential for this interaction. Next, in order to pinpoint residues that are important for RNase J multimerization, we constructed RNase J mutants in which nine charged residues of this C-terminal domain were individually replaced by alanine residues. As shown in Figure 3A, several mutants were significantly affected in their capacity to homo-oligomerize, including RNase J E618A, E654A, K660A, E663A and K649A. These data show that charged residues at the C-terminal domain of RNase J are important for its self-interaction.

**Figure 3.**
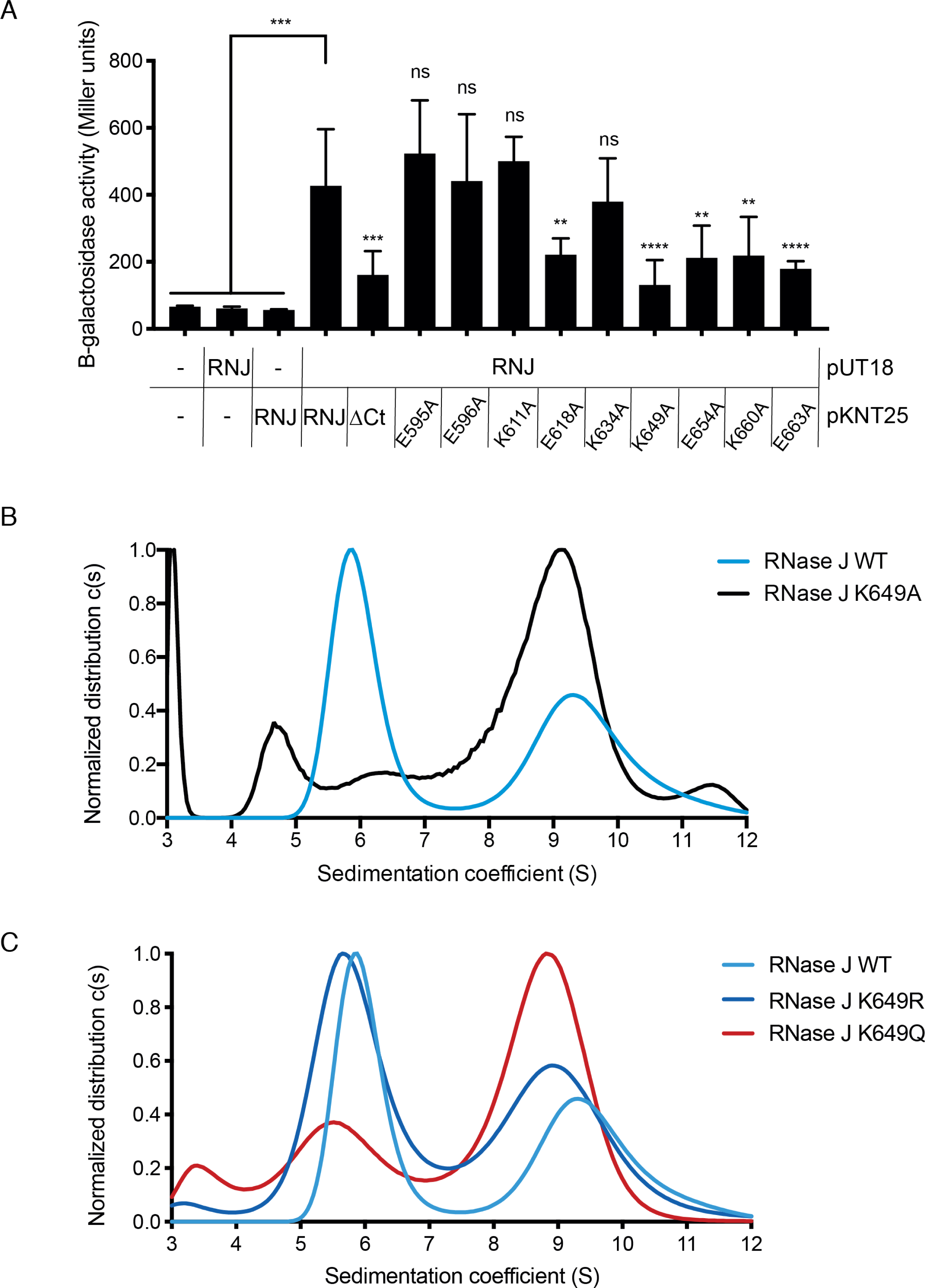
Point mutations modify the oligomeric state of RNase J. (A) BACTH two hybrid experiments to asses homo-oligomerization of WT and mutant RNase J proteins. Differences in the beta-galactosidase activity units are indicated with ns (non-significant) ** p-value<0.005, *** p-value<0.0005, **** p-value<0.0001. (B) Analytical ultracentrifugation sedimentation profiles of RNase J WT and variant K649A at 230 nm wavelength. (C) Analytical ultracentrifugation sedimentation profiles of RNase J WT and variants K649R and K649Q at 230 nm wavelength.

These residues are not at the predicted dimerization interface of the C-terminal domain, but they are likely to be involved in stabilizing the domain through interactions with side chains of complementary charge.

### RNase J is acetylated at several lysine residues

RNase R and RNase II from *E. coli* have been reported to be post-translationally modified by acetylation but no such modification was reported on RNase J. A mass spectrometry (MS) analysis was performed on an RNase J-FLAG fusion protein expressed from the *H. pylori* chromosome (Supp Figure S1) under the control of its native promoter. The RNase J-FLAG protein was pulled down from *H. pylori* crude lysates using anti-FLAG antibodies and extracted from a polyacrylamide gel. The experiment was performed on three biological replicates from cultures in exponential phase. We found that nine lysine residues of RNase J, spread across all protein domains, are consistently acetylated: K134, K140, K323, K337, K397, K511, K547, K634 and K649 (Supp Figure S2 and Supp Table S2).

### Lysine 649 is important for the oligomerization state of RNase J

Lysine 649 (K649), localized at the RNase J C-terminal domain, was particularly interesting since this residue was both acetylated and required for RNase J homo-oligomerization in two hybrid experiments. Therefore, AUC analyses were performed with purified RNase J variants (Supp Figure S1) to test the importance of K649 in RNase J homo-oligomerization. As shown in Figure 3B, the purified RNase J K649A mutant assembles exclusively as a dimer *in vitro*, with a complete loss of the monomeric population seen with the WT protein. Circular dichroism was performed to verify that this effect was not due to a defect in the folding of RNase J K649A. The circular dichroism spectra of wild-type and K649A variants of RNase J did not show significant differences (Supp Figure S3) and the estimated proportions of the different secondary structure elements were maintained (Supp Table S3). This shows that mutation K649A does not introduce a major folding defect in RNase J secondary structure. Thus, K649 is important for RNase J dimerization both *in vitro* and in *E. coli*.

### Acetylation mimicry of Lysine 649 affects the oligomeric state of RNase J

Given that K649 is acetylated, we next investigated whether RNase J acetylation at this residue could represent a control level of its oligomeric state. Therefore, we constructed RNase J mutants in which K649 was either replaced by an arginine residue (K649R), that mimics a non-acetylated state but maintains a positive charge at this position, or replaced by a glutamine residue (K649Q) which mimics an acetylated state while neutralizing the positive charge [a previously validated strategy^26^]. As shown in Figure 3C, the AUC sedimentation profile of RNase J K649R is similar to that of the wild type enzyme. In contrast, RNase J K649Q shows a significantly increased proportion of dimers over the monomers, similarly to the K649A mutant (Figure 3B-C). These data show that the positive charge of K649 is important for the stability of the RNase J monomeric state and that oligomerization is controlled by the charge state of K649.

### Mutations in K649 alter the exoribonuclease activity of RNase J

Since RNase J WT forms both monomers and dimers *in vitro*, we next assessed whether the oligomeric state and the presence of a charge at position 649 were correlated with differences in the *in vitro* activity of this protein. The exoribonuclease activity of RNase J was measured with a 24-nt long single stranded RNA substrate from the 5’-untranslated region (UTR) of *rnj*, upon which we previously found RNase J to be active^8^. This substrate was labeled with 6-carboxyfluorescein (6FAM) at either the 3’ or 5’-ends. As a control, a mutant inactivated at the RNase J active site, D299A, was constructed. We found, as expected, that the D299A inactive mutant did not degrade the substrate while the wild type RNase J was active in its 5’-3’ exoribonucleolytic mode (Figure 4A-B and Table 1). This activity is evident from the rapid depletion in the intact substrate bands labelled in 5’ (Figure 4A) or 3’ (Figure 4B) and, from the additional accumulation of the 1nt product upon digestion of the 5’-labeled substrate (Figure 4A).

**Figure 4.**
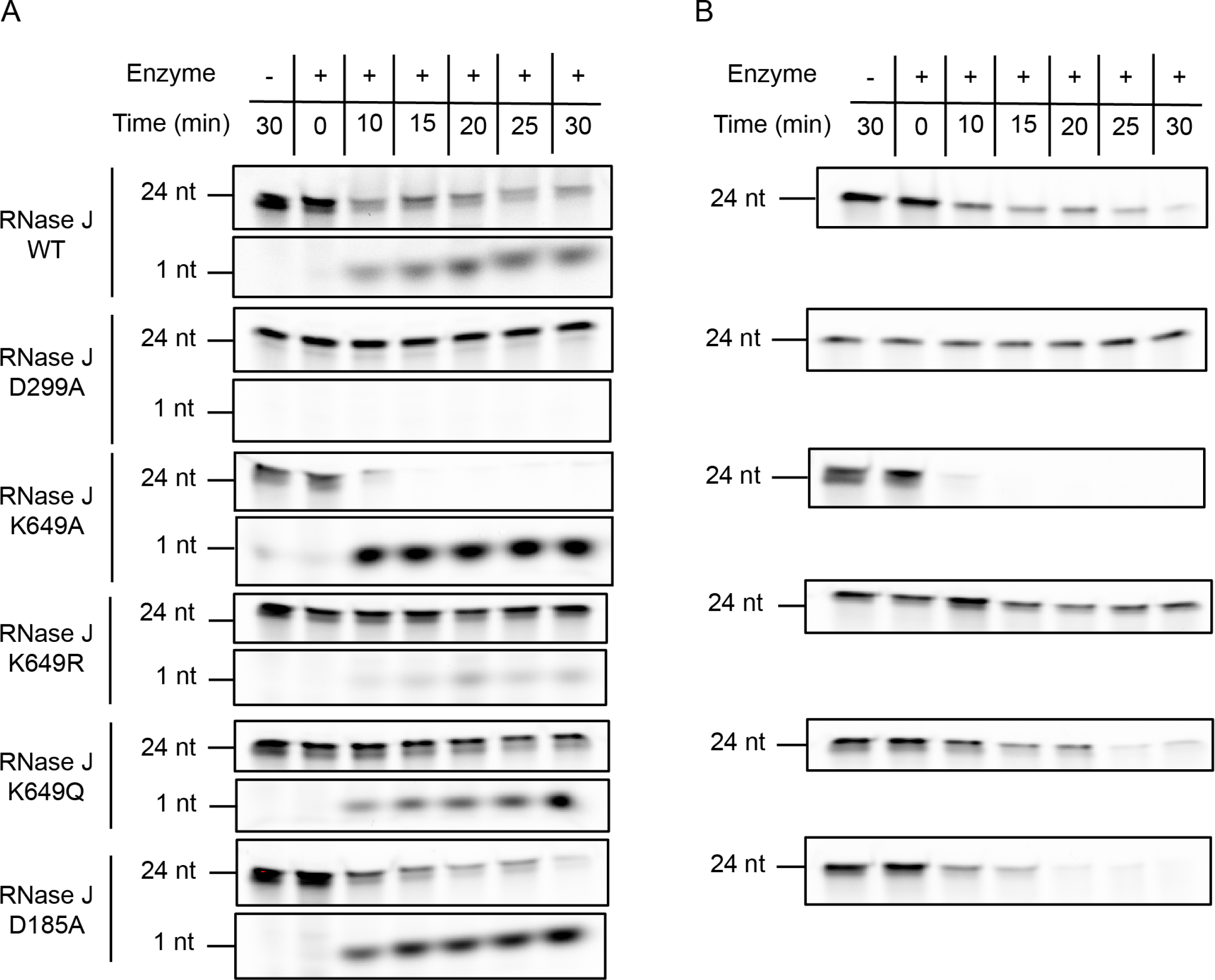
Exoribonuclease activity of RNase J wild type and mutants. (A) Exoribonuclease activity of RNase J WT and different variants using a 24-nt RNA substrate labeled with 6-carboxyfluorescein (6-FAM) at its 5’-end over the course of 30 min. (B) Exoribonuclease activity of RNase J WT and different variants using a 24-nt RNA substrate labeled with 6-FAM at its 3’-end over the course of 30 min.

The RNase J mutant with K649 replaced by R, that mimics a non-acetylated state, and which behaves as monomers and dimers like the WT protein, was strongly affected in its exoribonuclease activity (Figure 4 and Table 1). For the K649R variant, the half-lives of the 3’- and 5’-labeled substrates increased by 5 and 10-fold, respectively (Table 1).

In contrast, the protein variants with K649 replaced either by A (that abolishes both the positive charge at this position and the acetylation) or by Q (acetylation-mimicking mutation) and that both mainly assembled into dimers, presented an activity that was significantly higher than that of the RNase J K649R variant. The half-lives of 3’ and 5’-labeled substrates were 13 and 49-fold lower for the K649A variant and 2.5 and 4-fold lower for the K649Q variant, respectively. The half-lives of the substrates were calculated from three independent experiments for the wild type and for every RNase J variant and are shown in Table 1.

From these data, we conclude that, for these mutants, the dimeric form of RNase J is most probably the active form *in vitro*. Altogether, we identified a residue, K649, whose charge is important for RNase J activity and oligomerization and that is acetylated in *H. pylori* cells. This indicates a mechanism of control of RNase J activity that is based on the allosteric control of its dimerization by acetylation.

### The activity of the RNase J-RhpA complex is not affected by K649 mutations

It is not known whether the oligomerization state of RNase J can affect the formation of the RNA degradosome complex. Therefore, we wanted to determine whether mutations of K649 affect the activity of the RNase J-RhpA complex. For this, we used a 45-nt long RNA substrate with hairpin structures at both ends to prevent the exoribonuclease activity from the 5’-extremity and labeled it with 6FAM at the 3’ end [as in ^13^, Supp Figure S4A]. First, the activity of RNase J WT and mutant proteins alone was tested on this substrate. As compared to the linear 3’-labeled substrate, the degradation kinetics of this substrate by RNase J was significantly reduced but not abolished (half-life of the substrate is >4 fold lower, Table 1), suggesting that RNase J might attack this substrate by its endoribonuclease activity and then proceeds by exoribonuclease activity. As we previously reported, in a functional degradosome complex, RhpA will unwind the secondary structure elements and allow RNase J to act in its exoribonucleolytic mode. Most interestingly, the activity tests with a mixture of RNase J and RhpA in a 1:1 ratio resulted in a significant decrease in the half-life of the substrate to 6.3±2.9 min as compared to the activity of RNase J alone (approx. 31.4±17.6 min), compatible with a functional complex (Table 1). The half-life of the structured substrate incubated with the complex between RhpA and the RNase J K649A variant (that has increased exoribonuclease activity) was not affected (half-life 6.7±0.5 min) (Supp Figure S4B and Table 1). We conclude that the activity of the RNase J-RhpA complex is not significantly affected by the K649A mutation.

### Residue D185 is important for oligomeric state and the activity of RNase J

The RNase J structure suggests that residue D185 may indirectly support the stability of the monomer, as this residue is in an interaction network that supports a helical extension involving residues 584-594 that supports interactions with the C-terminal domain. In agreement with our predictions, AUC analysis of the RNase J D185A mutant revealed a much more predominant dimeric population, as compared to the wild type protein (Figure 5). Consistently with what we found for mutant K649A, this was correlated with higher exoribonucleolytic activity of RNase J D185A (Figure 4A-B and Table 1), suggesting again that the structural destabilization of the monomer leads to an increased amount of dimer with an enhanced enzymatic activity.

**Figure 5.**
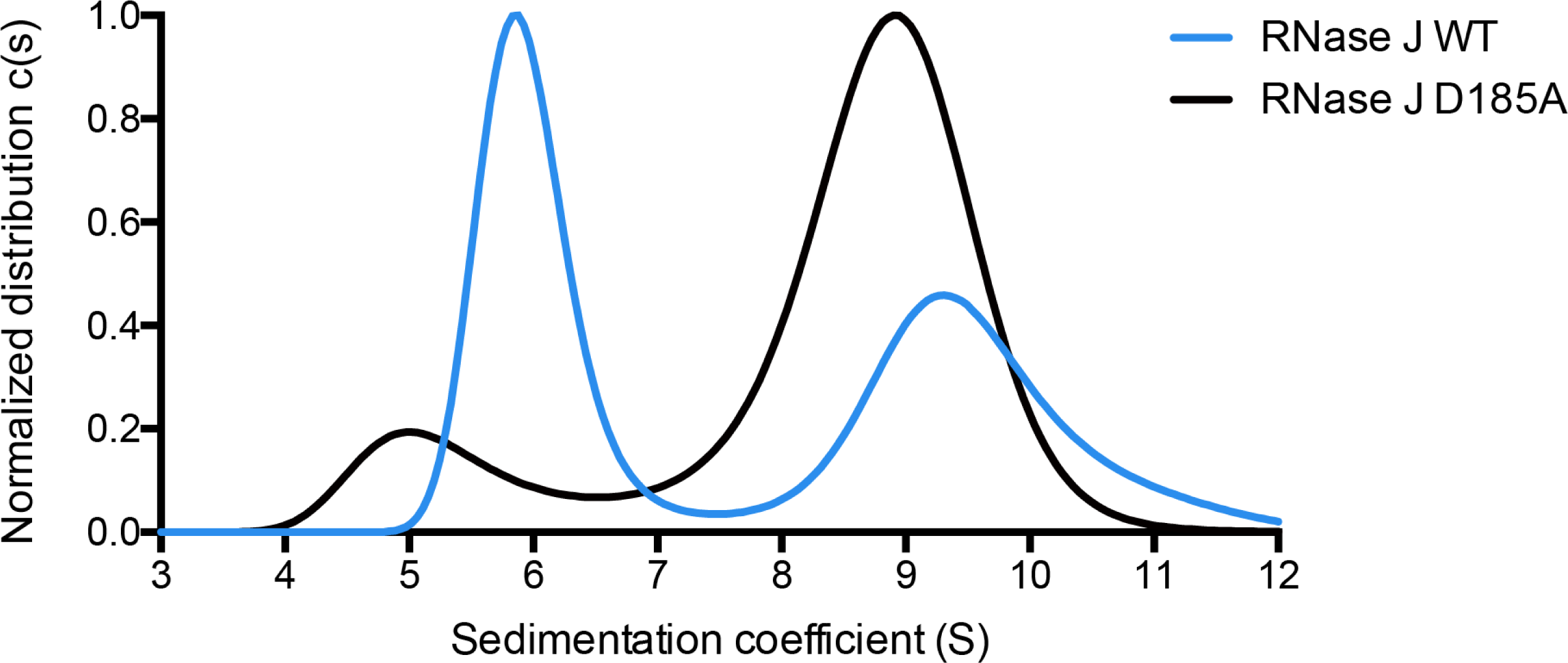
Impact of D185A mutation on RNase J oligomeric state. Analytical ultracentrifugation sedimentation profiles of RNase J WT and variant D185A, recorded at 230 nm wavelength.

### Analysis of the consequences of RNase J mutations on *H. pylori* morphology

We next wanted to examine the consequences of the mutations affecting RNase J oligomerization and/or acetylation state in live *H. pylori* cells. Since RNase J is an essential protein, strains carrying mutations in the acetylated residues identified above were constructed with the following strategy. *H. pylori* strain B128 was transformed with plasmid pILL2157 derivatives either expressing wild type or mutant variants of RNase J under the control of an IPTG-inducible promoter. Then, the chromosomal wild type copy of RNase J was deleted by homologous recombination and replacement with a kanamycin resistance cassette.

In *B. subtilis*, mutants deficient in RNase J1 present an altered cell morphology^27^. Therefore, the morphology of our *H. pylori* RNase J mutant strains was examined by phase contrast microscopy and quantified with the MicrobeJ ImageJ plug-in^28^. The wild type B128 strain presents cells that are relatively heterogeneous in shape and with an average length of 2.15±0.67 µm. We observed that, in a strain expressing wild type RNase J from the system described above, the *H. pylori* cells were longer in the absence of IPTG [lower amount of RNase J as compared to the WT strain^8^], than upon induction with 1 mM IPTG (where the expression of RNase J is induced). The cell length was 2.44±0.61 µm without IPTG and 2.05±0.51 µm with IPTG (Figure 6). This indicates that in *H. pylori*, RNase J is also regulating a factor required to maintain wild type cell morphology. Interestingly, we noted that the observed phenotype is not dependent on the activity of RNase J alone but rather on the RNA degradosome complex, as IPTG induction of RNase J WT in a strain lacking RhpA fails to complement this phenotype (average length of 3.41±0.83 µm) (Figure 6A).

**Figure 6.**
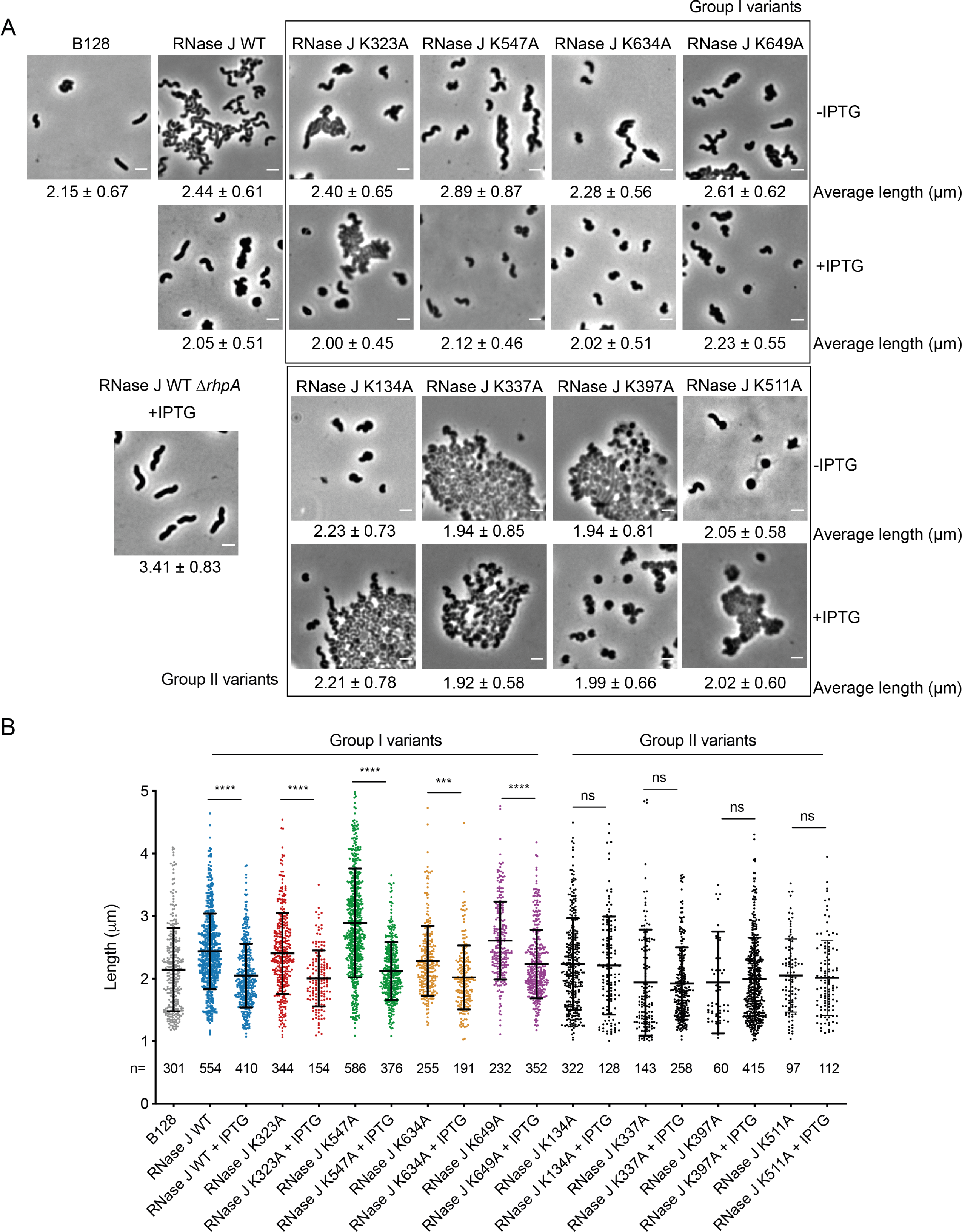
*Helicobacter pylori* cell morphology is affected by RNase J abundance and point mutations at lysine residues that are acetylation targets. (A) Phase contrast microscopy images of *H. pylori Δrnj* B128 strains expressing WT or mutant versions of RNase J from plasmid pILL2157 without (top panels) or with IPTG induction (bottom panels). The phenotype of a *H. pylori Δrnj*-*ΔrhpA* strains expressing WT RNase J from a plasmid is also shown. The scale bar represents 2 µm. (B) Dot plots showing the distribution of the cell lengths of wild type and the different mutants without or with IPTG. Horizontal lines indicate the mean length and the error bars are the standard deviation. ns (non-significant), ***p-value<0.0005, ****p-value<0.0001.

When we performed these tests with strains that expressed different RNase J variants, we identified two groups of mutations. In the first group, the RNase J variants behaved similarly to the wild type enzyme, as their expression upon IPTG addition resulted in a reduction of bacterial length. This was the case for RNase J K323A, K547A and K649A and, to a lesser extent, RNase J K634A. In the second group, that includes variants K134A, K140A, K337A, K397A and K511A, the strains presented a strong growth defect and substantial cell lysis and cellular debris were observed. In these cases, the cells were short with a tendency towards a coccoid morphology that is typical of *H. pylori* cultures under stress^29^. This phenotype was particularly evident in the case of cells expressing variant K397A. For these cultures, addition of IPTG favored bacterial growth but failed to restore the typical helical form of healthy *H. pylori* cells, suggesting that RNase J function is strongly impaired.

These results show that RNase J activity is important for the correct regulation of growth and cell morphology in *H. pylori* and that several of the residues that we found to be acetylated are important for RNase J function *in vivo*.

### RNase J is acetylated through a range of mechanisms in *H. pylori*

Several pathways contribute to acetylation in bacteria^30^. We tried to elucidate which are the enzymes responsible for the different acetylation events that occur on the RNase J of *H. pylori*. Therefore, we used the *H. pylori* strain expressing an RNase J-FLAG fusion from the chromosome to introduce deletions in genes that we predicted as acetylase candidates. The RNase J-FLAG protein was enriched from these strains and analyzed by MS as explained above. The tests were performed from three biological replicates. The acetylase candidate genes were *hpb8_615* [henceforth referred to as *rimI*, as it presents homology to *rimI* from *E. coli*^31^] and *hpb8_1270* [that contains two Gcn5-acetyltransferase (GNAT) domains]. Non-enzymatic acetylation by acetyl-CoA and acetyl-phosphate also occurs in bacteria^30, 32^. The impact of this pathway was assessed in *H. pylori* by constructing a mutant in the *pta-ackA* operon that is in charge of acetyl-phosphate synthesis and metabolism. As shown in Figure 7, we found that several pathways influence the acetylation of RNase J lysine residues. RimI is partially responsible for the acetylation of RNase J residues K323, K397 and K649. The HPB8_1270 protein is partially responsible for the acetylation of residues K323, K397, K511 and K649. Acetyl phosphate-mediated non-enzymatic acetylation participated in the acetylation of K134, K323, K397 and K511 and K649. For other residues, namely K140, K337, K547 and K634, no change in acetylation levels was detected in the mutants, suggesting that other acetylation mechanisms are likely at play in *H. pylori*. These results suggest a high redundancy of the different acetylation mechanisms that act on RNase J, particularly on residues K323, K397 and K649. Thus, multiple mechanisms participate in the acetylation of RNase J on different residues.

**Figure 7.**
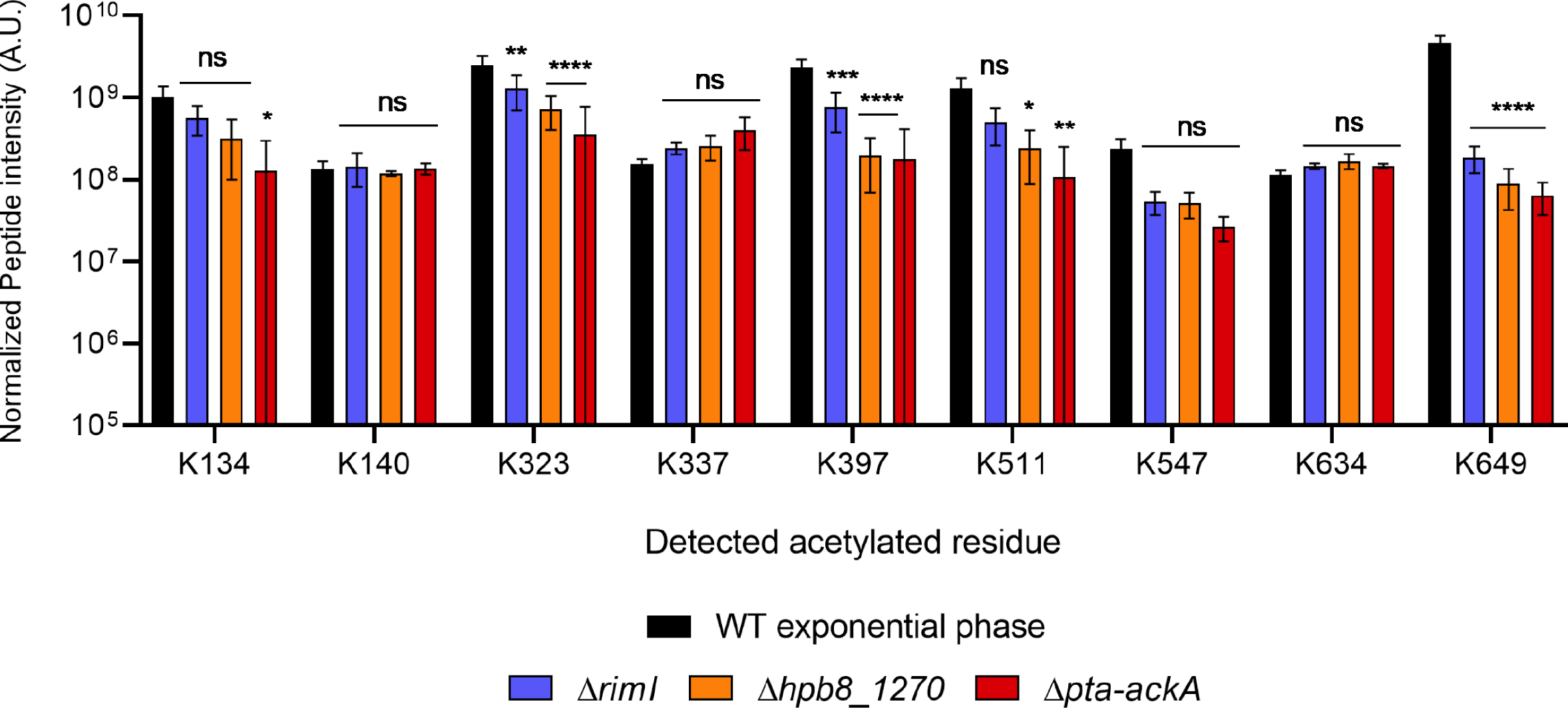
RNase J peptides containing acetylated residues extracted from different *H. pylori* strains. The normalized intensities were obtained by mass spectrometry from strains expressing FLAG-tagged RNase J in a wild-type background or in a mutant strain lacking different acetylation mechanism. ns (non-significant) p-value>0.05, *p-value<0.05, **p-value<0.005, ***p-value<0.0005, ****p-value<0.0001. Error bars represent the standard deviation.

## Discussion

In this study, we defined the structure of the major *H. pylori* ribonuclease RNase J and its *in vitro* oligomerization state alone and in the RNA degradosome with its RhpA RNA helicase partner. We also identify *in vivo* acetylation patterns of RNase J and demonstrate that these impact on oligomerization and activity *in vitro* and *in vivo*.

We found, by AUC, that RNase J is in equilibrium between monomeric and dimeric states. Earlier studies showed that *D. radiodurans* RNase J is also distributed between monomers and dimers^13^. *Bsu*RNase J1 forms dimers and tetramers *in vitro*^14^ and, when mixed with *Bsu*RNase J2, heterotetramers^10^. Similarly, RNase J1 from *S. aureus* forms homodimers and homotetramers *in vitro*^33^. RNase J of *S. pyogenes* crystallizes as tetramers that are composed of a dimer-of-dimers^12^. In the present study, at the studied *H. pylori* RNase J concentrations, no trace of tetramerization could be seen by AUC. It is likely that the observed tetrameric structure in the X-ray structure is due to crystal packaging and that it does not reflect the arrangement of RNase J in solution, as the dimer seems to be the active form. In addition, RhpA alone behaves as a monomer in solution by AUC. Together, RNase J and RhpA form a complex that is compatible with a 1:1 stoichiometry.

Acetylation has emerged in recent years as a major post-translational regulatory mechanism in prokaryotes^16^, and a few examples of its influence in RNases and RNA-degrading enzymes have been described, including *E. coli* RNase R^17^ and RNase II^18^ and *B. subtilis* CshA^19^. Here, we found that, in *H. pylori*, as many as 9 lysine residues of RNase J were consistently acetylated, although probably not at the same time, which constitutes the first evidence of protein acetylation in *H. pylori*. One of these residues is K649, which we also found to be involved in the oligomerization state of RNase J by BACTH in *E. coli*. Interestingly, this residue is located at the C-terminal domain of RNase J (Supp Figure S2). This region was also shown to be important for the oligomerization and exoribonucleolytic activity of the RNase J of *D. radiodurans*^13^. Therefore, we examined the role of K649 in RNase J oligomerization (by AUC) and exoribonuclease activity. Three mutants were analyzed, RNase J K649A that abolishes the side chain of the lysine residue, K649R that mimics a non-acetylated state while maintaining the positive charge and K649Q that mimics an acetylated state^26^. Whereas the K649R variant maintains a sedimentation profile similar to that of the wild type enzyme, variants lacking a positive charge in K649, either K649Q or K649A, had an increased proportion of dimer or displayed no monomeric form, respectively. Interestingly, we observed that the K649R variant is less active than the wild type RNase J whereas variants K649Q and K649A are more active than the K649R variant.

Our structural data indicated that residue K649 is not located at the interaction surface between the subunits of the dimer, but that it rather plays a structural role in the integrity of the overall structure. We conclude that elimination of the positive charge at residue 649 leads to a higher stabilization of the dimeric species. In agreement with this hypothesis, we observed that D185A, another mutation in a structurally important residue, had the same effect and favors the dimeric species. Interestingly, in *D. radiodurans*, the corresponding residue (D61) had already been identified as key for RNase J dimerization^13^. Electrostatic interactions of charged side chains on the surface of proteins can strongly contribute to the fold stability^23–25^. We propose that the network of surface electrostatic interactions stabilizes the CTD of RNase J, which in turn supports the homodimer. The CTD is the primary self-complementary interface of the dimer, and the interface between the core (amino acids 137-586) and between the dimer-of-dimers formed by the core both have marginal stability in isolation. Thus, changes in the charge, through acetylation, can thereby impact on stability of the CTD fold and oligomerization state.

The question naturally arises of how the catalytic activity of RNase J is affected by its oligomerization state. A clue is suggested in the overlay of the structure of *H. pylori* RNase J with the *S. coelicolor* homologue, in which there is a bound RNA substrate engaged at the active site (Pei et al., 2015) (Figure 1D). There are marked differences in the positions of loops at the entrance to the catalytic site that would obstruct the RNA from following the same route in the *H. pylori* enzyme, but the nucleic acid could approach the site from the opposite rim of the entrance. The exposed peptide loop regions are positioned to accommodate the CTD. It is possible that changes in the oligomerization state, through changes in CTD stability, will impact on the presentation of those loops and access to the active site in a gating mechanism.

Altogether, our results suggest that mutations that favor the dimeric RNase J conformation also increase the activity of RNase J of *H. pylori in vitro*. As the charge of residue 649 is important for dimerization, acetylation of K649 might tilt the balance of the RNase J species towards the dimeric form and potentially increase its activity. In this way, acetylation on K649 can modulate the activity of RNase J in *H. pylori* cells. Formation of the RNase J-RhpA complex is not affected by mutations or modifications of residue K649 in the ribonuclease, and thus such mutations do not significantly alter the activity of this complex. Earlier studies indicate that RNase J and the RNase J-RhpA complex can act on different transcripts^2, 8, 34^. It can be envisaged that acetylation of K649 would therefore have a differential impact on the subset of free RNase J that is not part of a degradosome assembly.

Furthermore, we addressed the role of the acetylated residues that we identified in *H. pylori* cells. It has been described that *B. subtilis* cells lacking RNase J1 show morphology defects with a tendency to filament^27^. We thus evaluated the morphology of *H. pylori* cells expressing RNase J variants, in which each acetylated lysine was replaced by an alanine residue. We observed that cells with a reduced concentration of wild type RNase J are longer than those in which RNase J expression is induced. This revealed that RNase J, most probably indirectly, is involved in the control of *H. pylori* length. Interestingly, this phenotype does not depend on the activity of RNase J alone, but rather on the RNA degradosome complex, since the overexpression of RNase J WT in a Δ*rhpA* strain does not reduce cell length. Given that we previously showed that about 80% of *H. pylori* mRNAs are at least two times more abundant in an RNase J-depleted mutant, it is difficult to define the target mRNA that is causing the observed elongation phenotype^8^. One possibility would be the overexpression of MreC (9.3-fold upregulated in an RNase J-depleted strain), that has been shown to cause cell filamentation in *H. pylori*^35^.

Alanine replacement of the different acetylated positions of RNase J leads to different phenotypes, showing that acetylation is indeed important for RNase J activity in *H. pylori*. Variant K649A behaves like the wild type enzyme in this test, in agreement with the notion that the morphology phenotype depends on the activity of the whole RNA degradosome complex, which is not affected by whether there is a charge in this position or not. The phenotype of the K134A mutant suggests that it is affected in RNase J activity; interestingly this residue is located within the N-terminal extension of *Hp*RNase J (Supp Figure S2), that we previously found to be very important for efficient RNase J activity^8^. However, it is not known how these acetylated residues impact RNase J properties. They could affect its stability, activity, oligomerization, degradosome complex formation, compartmentalization, binding of other partners or RNA, or a combination of these.

We found that several mechanisms participate in the acetylation of the different residues of RNase J. Our results indicate that HPB8_1270 and the *H. pylori* RimI homologue indeed function as acetylases. In addition, we uncovered a role of non-enzymatic acetylation by acetyl phosphate that is produced and metabolized by the Pta and AckA enzymes. The *E. coli* RimI protein is a multifunctional acetyltransferase, that can perform ε-lysine acetylation and Nα acetylation on the L31 and S18 ribosomal proteins, respectively^31, 36^, with high substrate specificity. In contrast, RimI from *Mycobacterium tuberculosis* is an Nα acetylase with more relaxed substrate specificity *in vitro*^37^. Here, we found that the *H. pylori* RimI targets K323, K397 and K649 of RNase J. HPB8_1270 is a protein belonging to the DUF2156 family, containing two GNAT domains. It presents homology with the BT_3689 protein from *Bacteroides thetaiotaomicron*, which contains an acetyl-CoA molecule in its active site (PDB 2HQY), suggesting its function as an acetyltransferase. Here, we found that HPB8_1270 is involved in the acetylation of K323, K397, K511 and K649, showing that in *H. pylori* this protein is most likely an acetyltransferase. Lastly, acetyl phosphate-mediated acetylation, which is responsible for most acetylation events in *E. coli* and *B. subtilis*^30, 32^, was important for the acetylation of K134, K323, K397, K511 and K649.

These data demonstrate that multiple mechanisms can target the same lysine residue and that there is a high redundancy in the functions of these acetylation mechanisms. Acetylation at residue K649 was strongly dependent on three such mechanisms; this redundancy could testify of a critical importance of acetylation at this position. In addition, other acetylation mechanisms could occur in *H. pylori* that might account for some of these acetylation events. More work is needed, in particular using multiple mutants to determine how many of these mechanisms can target the different residues and the influence of the different acetylation mechanisms on the activity and regulation of RNase J and the RNA degradosome.

Using Clustal Omega 1.2.4^38^, we performed an alignment of the RNase J proteins of *H. pylori* and of the closely related organism *Campylobacter jejuni*, but also of the more distant *B. subtilis*, *S. aureus*, *S. pyogenes* and *D. radiodurans*. Interestingly, this revealed that most of the acetylated lysines are conserved in *H. pylori* and *C. jejuni* RNase J proteins, and some of them are also conserved in more distant organisms (Supp Figure S5). The case of K511 is particularly interesting, as it is conserved in *S. pyogenes*, *B. subtilis*, *H. pylori* and *C. jejuni*, and in the case of *S. aureus* this position is a Q residue which might behave as a “permanently-acetylated” lysine. In *H. pylori*, the mutant expressing RNase J K511A variant presents a strong growth defect. The relevance of lysine acetylation in the regulation of the activity of the RNase J proteins from other organisms remains to be addressed.

In conclusion, we have established that RNase J is acetylated on multiple lysine residues through different mechanisms, and that its acetylated residues affect the activity of RNase J and phenotypes such as cell morphology. Furthermore, our data suggests that acetylation on K649 can influence the oligomeric state and activity of RNase J, and that the dimeric form could be the most active one. Thus, acetylation is an important player in the modulation of the activity of RNase J and of the RNA degradosome of *H. pylori*.

## Materials and methods

### Bacterial strains and growth conditions

The bacterial strains used in this study are summarized in Supp Table S4. *H. pylori* strains were derivatives of strain B128^39, 40^. Plasmids (Supp Table S4) were constructed and amplified using *E. coli* XL1-Blue (Agilent Technologies). *H. pylori* strains were grown on Blood agar base 2 (Oxoid) plates supplemented with 10% defibrinated horse blood and with an antibiotic-antifungal cocktail composed of 2.5 µg/mL amphotericin B, 0.31 µg/mL polymyxin B, 6.25 µg/mL trimethoprim and 12.5 µg/mL vancomycin. *H. pylori* mutants were selected using 20 µg/mL kanamycin, 10 µg/mL apramycin or 6 µg/mL chloramphenicol. For liquid cultures, we used Brucella broth supplemented with 10% fetal calf serum (FCS) (Eurobio), the antibiotic-antifungal cocktail and, when necessary, chloramphenicol to select for plasmid-containing bacteria. *H. pylori* cells were grown at 37°C under a microaerophilic atmosphere (6% O_2_, 10% CO_2_, 84% N_2_) using an Anoxomat (Mart Microbiology) atmosphere generator.

### Plasmid preparation and transformations

A NucleoBond Xtra midi kit (Macherey-Nagel) and a QIAamp DNA minikit (Qiagen) were used for plasmid preparations for *H. pylori* transformation and genomic DNA extractions, respectively. For plasmid preparations for *E. coli* transformation and sequencing, a QIAprep Spin Miniprep kit (Qiagen) was used. PCR was carried out with either DreamTaq DNA polymerase (Thermofisher) or with Q5 High-Fidelity DNA polymerase (NEB) when the product required a high-fidelity polymerase. A list of the oligonucleotides that were used can be found in Supp Table S5.

### Construction of H. pylori mutants and fusions

The cassette for the construction of the *H. pylori* RNase J-FLAG strain was constructed by the isothermal assembly technique^41^, by assembling in frame the last 500 bp of the *rnj* gene followed by the FLAG-tag, the apramycin resistance gene and the 500 bp immediately downstream from *rnj*. The resulting fragment was PCR-amplified and 1 µg of the product was introduced into *H. pylori* by natural transformation. Transformants were selected on plates with apramycin. Strains carrying the pILL2157 plasmid with the wild type *rnj* gene^2^ and its derivates with the different *rnj* mutants (constructed following standard procedures with the primers compiled in Supp Table S5) were transformed in the same manner with 1.5 µg of the plasmids and selected on plates with chloramphenicol. The resulting strains were then used to transform by cassettes constructed in a similar manner to delete the *rnj* gene and replace it by a kanamycin resistance gene [as in ^2^]. The transformants were selected on plates containing kanamycin, chloramphenicol and 1 mM IPTG to allow the expression of the copy of *rnj* from the plasmid.

Plasmids for BACTH were constructed from plasmids pUT18 and pKNT25^42^ by inserting wild type or mutant variants of the *rnj* gene in the *Pst*I and *Kpn*I restriction sites following standard techniques using plasmids indicated in Supp Table S4.

Deletion of the genes of interest, construction of fusions and/or insertion of cassettes was verified by PCR and sequencing of the region of interest.

### Purification and crystallography of RNase J

A vector co-expressing residues 21-465 of RhpA with a C-terminal 3C-HALO-his6 tag and residues 139-691 of RNase J (C1hRSEj) (from *H. pylori* strain 26695 gene HP1430, UniProt P56185) with a C-terminal hexa-histidine tag was generated by the Oxford Protein Production Facility at Harwell. Cells were grown in terrific broth supplemented with 0.5 M NaCl and 1 mM betain, 30 µg/mL kanamycin, 50 µg/mL carbenicillin and 20 µg/mL chloramphenicol. Cells were grown at 37°C until optical density at 600 nm reached 0.3, then moved to 20°C and induced with 0.2 mM IPTG (isopropyl β-D-1-thiogalactopyranoside) for 24 hours. Cells were harvested by centrifugation, then suspended in 50 mM Tris pH 7.4, 50 mM HEPES (4-(2-hydroxyethyl)-1-piperazineethanesulfonic acid) pH 7.4, 500 mM NaCl, 1 mM TCEP (tris(2-carboxyethyl)phosphine), 0.05% v/v Triton X-100, 5% v/v glycerol and 10 mM imidazole and stored frozen. The suspension was thawed, passed 5 times through a cell disruptor (Avestin Emulsiflex C5 homogeniser, 1000 bars) and clarified by centrifugation at 37,000 g for 30 minutes, 5°C. The supernatant was applied to a HiTrap Ni chelating column, washed with lysis buffer containing 0.01% v/v Triton X-100, and 15 mM imidazole, then eluted with a gradient in imidazole. The optimal fractions were pooled and cleaved with 3CP-GST protease cleavage, using 300 µl of 0.85 mg/mL protease to cleave C1/RSE protein isolated from 1 L culture, at 4°C overnight. This step removes HALO-Chis-tag on RhpA. The 3CP-cleaved protein was loaded onto HALO-RESIN and the flow-through collected, which contained the RNase J/helicase complex. The sample was concentrated with a 30kD MWCO centrifugal concentrator and applied to a S200 SEC column equilibrated in 20 mM Tris-HCl pH 7.4, 20 mM HEPES pH 7.4, 150 mM NaCl, 5% glycerol, 0.1% v/v Triton X-100, 1 mm TCEP. The sample was concentrated with a 30kD MWCO centrifugal concentrator to optical density at 280 nm of 3.38 and used in crystallization screens. Crystals were found in PEG I screen 0.1 M HEPES pH 7.5, 25% wt/v polyethylene glycol 1000.

Diffraction data were collected at I24 at Diamond Light source at wavelength 0.9778 Å using a Pilatus6M detector. Crystals are in space group P4_1_22 with cell dimensions a=b=158 c=214.26 Å. The structure was solved by molecular replacement using *Streptomyces* RNase J (PDB 3BK2) for with PHASER and the CCP4 suite^43, 44^. There was no space in the lattice for the RhpA protein. The model of RNase J 139-691 was built using COOT^45^ and refined with PHENIX^46^. A homology model for the C-terminal domain, residues 587 to 691, was prepared using PHYRE2^47^, and the structure of the full-length protein was predicted using the ROBETTA server^21^. The C-terminal domain from the ROBETTA model was docked into the map and fitted as a rigid body and then linked to the model of the well resolved core (139-586). As the density for the C-terminal domain was poorly resolved due to disorder, the side chains were removed. A final cycle of refinement using the core only (137-586) and not the disordered C-terminal domain were used to provide the refinement parameters summarized in the supplementary Table S1. A monomer of RNase J 139-691 occupies the asymmetric unit, and the tetramer is generated through crystallographic symmetry. Estimates of buried surface area were made using the PISA program in the CCP4 suite^48^. The model and structure factors have been deposited in the PDB with accession code 7PCR.

### Bacterial two-hybrid (BACTH)

Bacterial two hybrid assays were carried out in *E. coli* strain BTH101 as previously described^42^. Briefly, strains carrying derivatives of the two vectors, pUT18 and pKNT25, were grown overnight in 1 mL LB with 40 µg/mL kanamycin, 100 µg/mL ampicillin and 0.1 mM IPTG. The OD at 600 nm of the resulting cultures was measured in a TECAN plate reader and the beta-galactosidase activity was calculated by mixing 500 µL of the cultures with 500 µL of buffer Z, 100 µL of chloramphenicol and 50 µL of 0.1% SDS, vortexing the cells and then adding 200 µL of 4 mg/mL ONPG. The reactions were incubated at 28°C until they turned yellow and the reactions were stopped by adding 500 µL of 1 M Na_2_CO_3_. Samples were centrifuged for 5 min at 14,000 rpm and the OD at 420 and 550 nm of the upper fraction was measured in a TECAN plate reader. The beta-galactosidase activity (Miller units) was calculated as described^49^.

### Protein expression and purification

Wild type and variant versions of RNase J and RhpA were cloned into a pET28a+ vector in *E. coli* XL1-Blue and then transformed in *E. coli* Bli5 for expression. The strains were grown in 800 mL cultures in LB with 40 µg/mL kanamycin and 25 µg/mL chloramphenicol and, once they reached an OD of 0.5-0.8, were induced with 1 mM IPTG. The cultures were incubated at 30°C and 160 rpm during 3 h. The cells were collected by centrifugation at 4000 rpm in an Avanti J-E centrifuge (Beckman Coulter) with a JA-14 rotor at 4°C during 30 min and pellets were frozen at -80°C until further processing.

Pellets were thawed on ice and resuspended on lysis buffer containing 25 mM Tris-HCl pH 7.8 for RNase J and pH 8 for RhpA, 500 mM NaCl, 10% glycerol, 20 mM imidazole, 1 mM dithiothreitol (DTT), 50 U benzonase and protease inhibitor cocktail (c0mplete EDTA-free, Roche). Cells were lysed by sonication and centrifuged for 30 min at 4,000 rpm in a 5810R centrifuge (Eppendorf) at 4°C to eliminate cellular debris. The supernatant was then incubated overnight at 4°C and under agitation with 2 mL Ni-NTA resin (50% v/v, Thermofisher) pre-equilibrated with lysis buffer.

Resins were packed onto Poly-Prep Chromatography columns (Bio-Rad) and washed with 10 volumes of 25 mM Tris-HCl at the appropriate pH, 300 mM NaCl, 1 mM DTT and 20 mM imidazole, 10 volumes of the same buffer with 50 mM imidazole and 10 volumes of the same buffer with 100 mM imidazole. The elution was carried out with an elution buffer containing 25 mM Tris-HCl at the appropriate pH, 300 mM NaCl, 1 mM DTT and 500 mM imidazole, in 1 mL fractions. Aliquots of the resulting fractions were run on a 4-20% Mini-Protean TGX stain-free precast protein gel (Bio-Rad) at 200V for 30 min and stained by Imperial protein stain (ThermoFisher). After destaining, the corresponding purest fractions were chosen, mixed and concentrated using Vivaspin 20 tubes (MWCO 30,000 Da, GE Healthcare) down to 1 mL volumes. The buffer was replaced using G-25 columns (PD MidiTrap G-25, GE Healthcare) with 25 mM Tris-HCl, 300 mM NaCl, 1 mM DTT and 10% glycerol. Samples were aliquoted and frozen at -80°C.

### Analytical ultracentrifugation

Samples of purified proteins were thawed on ice and the buffer was replaced using G-25 columns (PD MiniTrap G-25, GE Healthcare) with 25 mM Tris-HCl pH 8, 300 mM NaCl and 0.5 mM Tris(2-carboxyethyl)phosphine (TCEP, Sigma).

Appropriate amounts of the different proteins were mixed and diluted in the same buffer with a final volume of 300 µL that was charged into 2 sector analytical ultracentrifugation cells. The concentration of the proteins was 3 mM when there was only one protein, 3 mM:3 mM when two proteins were in equimolar amounts and 1 mM:5 mM when the proteins were mixed at a 1:5 ratio. The centrifugation was performed for 16 h on an Optima analytical ultracentrifuge (Beckman-Coulter) with an 8-hole AnTi50 rotor, at 42,000 rpm and 20°C, measuring the optical density at 230 nm in order to determine the sedimentation profile of the protein or complex. The data was analyzed with the Sedfit software (NIH) using a diffusion deconvoluted continuous sedimentation coefficient distribution c(s) with one discrete component model^50^. The model was chosen in order to take in consideration the signal due to TCEP absorption. The presented graphs correspond to a resolution of 300 points with a confidence interval of 0.9. For the individual proteins, the sedimentation coefficient independent of the concentration (S_0_) was calculated.

The partial specific volume (*v̄*) of the proteins was estimated by the software Sednterp^51^ based on the amino acid sequence. This software was also used for the calculation of the density and viscosity of the buffer used throughout the experiments. Calculation of sedimentation coefficients for protein structures was performed with the program Hydropro^52^.

### Circular dichroism

The near-UV absorption spectra (195-260 nm) were measured, in the same buffer as for the AUC experiments, by circular dichroism with a Circular Dichroism Spectrometer Model 215 (Aviv instruments) using 0.2 mm optical path cells with 5 mM of RNase J WT and K649A mutant. The proportions of the different secondary structure elements were determined with the BeStSel software ^53, 54^ based on the amino acid sequence.

### Ribonuclease activity tests

The endoribonuclease and exoribonuclease activities were measured using 6-carboxyfluorescein (6FAM)-labeled RNA substrates either in 5’ or in 3’ as listed in Supp Table S5. 500 nM RNA substrate were incubated with 500 nM protein in a total volume of 10 µL with a buffer containing 50 mM Tris-HCl pH 7.8, 100 mM NaCl, 8 mM MgCl_2_, 0.1 mM and 0.1 mg/mL bovine serum albumin (BSA). The reactions were carried out at 37°C for 0-30 min for the exoribonuclease activity tests and 0-75 min for endoribonuclease activity tests. Reactions were stopped by adding 98% formamide and 10 mM ethylene diamine tetraacetic acid (EDTA) and by incubating for 15 min at 95 °C.

The degradation products were run on a Mini-Protean TBE-Urea gels (Bio-Rad) that was pre-run for 60 min at 200 V. The samples were run for 5 min at 50 V and then for 30 min at 200 V. Gels were visualized on a Chemidoc MP Imaging System (Bio-Rad) with 0.2 s exposure at 488 nm. The gels were then stained with ethidium bromide to reveal the molecular weight marker (ssRNA molecular weight marker, Takara). Experiments were performed in triplicate and images were quantified ImageLab (Bio-Rad, version 6.1). Decay coefficients, λ, were calculated with the formula *N(t) = N_o_e^-At^*, where *N(t)* is the amount of substrate at time *t* and *N_0_* is the initial amount. The half-lives, *t_1/2_* were then calculated with the formula 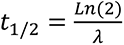.

### Phase contrast microscopy

Phase contrast microscopy was performed with an Axio Observer microscope (Zeiss) equipped with an Axiocam camera with X100 magnification. Image acquisition was performed with the axiovision software. Images were cropped and adjusted using FiJi software. The length of wild type cells and that of cells expressing wild type or mutant versions of RNase J from a plasmid, with or without 1 mM IPTG, was measured by using the MicrobeJ plugin of ImageJ ^28^ with the following parameters: cell area above 0.7 µm^2^, minimal length of 1 µm and cell width between 0.4-1 µm.

### Pulldown and mass spectrometry

*H. pylori* cultures of strains expressing FLAG-tagged RNase J were grown in 15 mL of Brucella medium until an OD_600 nm_ of 0.6-1.2 for exponential phase cultures. Cells were pelleted by centrifugation for 10 min at 4 °C and 4,000 rpm and resuspended in PBS to an OD_600 nm_ of 10. Cells were lysed by sonication and cell debris were eliminated by centrifugation for 10 min at 13,000 rpm and 4 °C. The supernatant constitutes the total extract fraction.

Pulldowns were performed using 2 µL of monoclonal anti-FLAG M2 antibodies produced in mice (F3165, Sigma) per reaction and 50 µL of magnetic protein G-coated Dynabeads according to the supplier’s instructions (Dynabeads Protein G for Immunoprecipitation, 10003D, ThermoFisher). Proteins were eluted in the denaturing mode and run in a 4-20% Mini-Protean TGX stain-free precast protein gel (Bio-Rad) at 200V for 30 min and stained by Imperial protein stain (ThermoFisher) and destained.

Each band corresponding to RNase J-FLAG was cut and washed several times in 50 mM ammonium bicarbonate (ABC)-acetonitrile (ACN) (1:1) for 15 min at 37 °C. Disulfide bonds were reduced with 10 mM DTT and cysteines alkylated with 55 mM chloroacetamide. Trypsin (V5111, Promega) digestion was performed overnight at 37 °C in 50 mM ABC. Peptides were extracted from the gel by two incubations in 50 mM ABC-ACN-formic acid (FA) (50:50:0.5) for 15 min at 37 °C. After ACN evaporation in a Speed-Vac, resulting peptides were desalted with stage-tip ^55^ using C18 Empore disc and eluted with 80% ACN, 0.1% FA. Peptides were resuspended in 2% ACN, 0.1% FA prior to liquid chromatography coupled to mass spectrometry (LC-MS/MS) analysis.

For LC-MS/MS, a nanochromatographic system (Proxeon EASY-nLC 1200 - ThermoFisher Scientific) was coupled on-line to a Q Exactive^TM^ Plus Mass Spectrometer (ThermoFisher Scientific) using an integrated column oven (PRSO-V1 - Sonation GmbH, Biberach, Germany). For each sample, peptides were loaded on an in-house packed 26 cm nano-HPLC column (75 μm inner diameter) with C18 resin (1.9 μm particles, 100 Å pore size, Reprosil-Pur Basic C18-HD resin, Dr. Maisch GmbH, Ammerbuch-Entringen, Germany) after an equilibration step in 100 % solvent A (H_2_O, 0.1% FA).

Peptides were eluted with a multi-step gradient using 2 to 7 % solvent B (80% ACN, 0.1% FA) during 5 min, 7 to 23% during 70 min, 23 to 45% during 30 min and 45 to 95% during 5 min at a flow rate of 300 nL/min over 132 min. Column temperature was set to 60°C.

MS data were acquired using Xcalibur software using a data-dependent Top 10 method with a survey scans (300-1700 m/z) at a resolution of 70,000 and a MS/MS scans (fixed first mass 100 m/z) at a resolution of 17,500. The AGC target and maximum injection time for the survey scans and the MS/MS scans were set to 3E6, 20 ms and 1E6, 60 ms respectively. The isolation window was set to 1.6 m/z and normalized collision energy fixed to 28 for HCD fragmentation. We used a minimum AGC target of 1E4 for an intensity threshold of 1.7E5. Unassigned precursor ion charge states as well as 1, 7, 8 and >8 charged states were rejected and peptide match was disable. Exclude isotopes was enabled and selected ions were dynamically excluded for 30 seconds.

Raw data were analyzed using MaxQuant software version 1.6.6.0^56^ using the Andromeda search engine^57^. The MS/MS spectra were searched against a UniProt *Helicobacter pylori* [strain B8^39, 40^] database (download in 24/06/2019, 1,719 entries) and tagged RNase J. Usual known mass spectrometry contaminants and reversed sequences of all entries were included.

Andromeda searches were performed choosing trypsin as specific enzyme with a maximum number of five missed cleavages. Possible modifications included carbamidomethylation (Cys, fixed), oxidation (Met, variable), Nter acetylation (variable) and K acetylation (Lys, variable) for in-gel peptides. The mass tolerance in MS was set to 20 ppm for the first search then 4.5 ppm for the main search and 20 ppm for the MS/MS. Maximum peptide charge was set to seven and five amino acids were required as minimum peptide length.

A false discovery rate (FDR) cutoff of 1 % was applied at the peptide and protein levels.

Raw files and MaxQuant files (msms.txt) were processed in Skyline-daily 20.1.9.268^58^ to generate MS1-XIC and perform peak integration for each K-acetylated peptide. Normalization across samples was performed by using the mean of the ratios from non-modified peptides (Supp Table S2).

The mass spectrometry data have been deposited to the ProteomeXchange Consortium via de PRIDE partner repository^59^ with the dataset identifier PXD027670.

### Statistical analyses

Statistical analysis was performed using Prism 9. We applied one-way ANOVA with Dunnett’s multiple comparisons test in Figure 2A, one-way ANOVA with Sidak’s multiple comparisons test in Figure 6B and two-way ANOVA with Dunnett’s multiple comparisons test in Figure 7.

## Acknowledgements

We would like to thank Sébastien Brûlé and Bruno Baron for their help with AUC and circular dichroism experiments, and Frédéric Fischer for bioinformatical analysis regarding protein HPB8_1270. We thank Diane Soussan for her help in preliminary experiments of this project and the members of the Unité Pathogenèse de *Helicobacter*. ATA is part of the Pasteur - Paris University (PPU) International PhD Program. This project has received funding from the European Union’s Horizon 2020 research and innovation program under the Marie Sklodowska-Curie grant agreement N° 665807, and from the Institut Carnot Pasteur Microbes & Santé. MBL was a master student of the UTC (Université de Technologie de Compiègne). Support was provided by “Fondation pour la Recherche Médicale” for the grant DBF20161136767 to HDR and the Pasteur-Weizmann Consortium of “The Roles of Noncoding RNAs in Regulation of Microbial Life Styles and Virulence” to HDR. BFL and XYP are supported by a Wellcome Trust Investigator award (200873/Z/16/Z). The authors would like to thank the DIM 1HEALTH region Ile-de-France for funding the Centrifection project that has allowed the Optima ultracentrifuge investment. We thank the staff of the OPPL at Harwell Campus, UK for preparing vectors, screening expression conditions for the recombinant proteins, and helpful discussions.

## References

1. Tejada-Arranz, A., de Crécy-Lagard, V. & de Reuse, H. Bacterial RNA Degradosomes: Molecular Machines under Tight Control. Trends Biochem. Sci. 45, 42–57 (2020).

2. Redko, Y. et al. A minimal bacterial RNase J-based degradosome is associated with translating ribosomes. Nucleic Acids Res. 41, 288–301 (2013).

3. Amieva, M. & Peek, R. M. Pathobiology of *Helicobacter pylori*–Induced Gastric Cancer. Gastroenterology 150, 64–78 (2016).

4. Sharma, C. M. et al. The primary transcriptome of the major human pathogen Helicobacter pylori. Nature 464, 250–255 (2010).

5. Eisenbart, S. K. et al. A Repeat-Associated Small RNA Controls the Major Virulence Factors of *Helicobacter pylori*. Mol. Cell 80, 210–226 (2020).

6. Tejada-Arranz, A. & De Reuse, H. Riboregulation in the Major Gastric Pathogen *Helicobacter pylori*. Front. Microbiol. 12, 712804 (2021).

7. El Mortaji, L., et al. The sole DEAD-box RNA helicase of the gastric pathogen *Helicobacter pylori* is essential for colonization. MBio 9, e02071–17 (2018).

8. Redko, Y. et al. RNase J depletion leads to massive changes in mRNA abundance in *Helicobacter pylori*. RNA Biol. 13, 243–253 (2016).

9. Tejada-Arranz, A. et al. The RNase J-Based RNA Degradosome Is Compartmentalized in the Gastric Pathogen *Helicobacter pylori*. MBio 11, e01173–20 (2020).

10. Mathy, N. et al. *Bacillus subtilis* ribonucleases J1 and J2 form a complex with altered enzyme behaviour. Mol. Microbiol. 75, 489–498 (2010).

11. Hausmann, S. et al. Both exo- and endo-nucleolytic activities of RNase J1 from *Staphylococcus aureus* are manganese dependent and active on triphosphorylated 5′-ends. RNA Biol. 14, 1431–1443 (2017).

12. Pei, X.-Y., Bralley, P., Jones, G. H. & Luisi, B. F. Linkage of catalysis and 5’ end recognition in ribonuclease RNase J. Nucleic Acids Res. 43, 8066– 76 (2015).

13. Zhao, Y. et al. Structural insights into catalysis and dimerization enhanced exonuclease activity of RNase J. Nucleic Acids Res. 43, 5550–5559 (2015).

14. Newman, J. A. et al. Unusual, Dual Endo- and Exonuclease Activity in the Degradosome Explained by Crystal Structure Analysis of RNase J1. Structure 19, 1241–1251 (2011).

15. Wolfe, A. J. Bacterial protein acetylation: new discoveries unanswered questions. Curr. Genet. 62, 335–341 (2016).

16. Carabetta, V. J. & Cristea, I. M. Regulation, Function, and Detection of Protein Acetylation in Bacteria. J. Bacteriol. 199, e00107–17 (2017).

17. Liang, W., Malhotra, A. & Deutscher, M. P. Acetylation Regulates the Stability of a Bacterial Protein: Growth Stage-Dependent Modification of RNase R. Mol. Cell 44, 160–166 (2011).

18. Song, L., Wang, G., Malhotra, A., Deutscher, M. P. & Liang, W. Reversible acetylation on Lys501 regulates the activity of RNase II. Nucleic Acids Res. 44, 1979–1988 (2016).

19. Ogura, M. & Asai, K. Glucose Induces ECF Sigma Factor Genes, sigX and sigM, Independent of Cognate Anti-sigma Factors through Acetylation of CshA in Bacillus subtilis. Front. Microbiol. 7, 1918 (2016).

20. Callebaut, I., Moshous, D., Mornon, J. P. & De Villartay, J. P. Metallo-β-lactamase fold within nucleic acids processing enzymes: The β-CASP family. Nucleic Acids Res. 30, 3592–3601 (2002).

21. Baek, M. et al. Accurate prediction of protein structures and interactions using a three-track neural network. Science 373, 871–876 (2021).

22. Nooren, I. M. A. & Thornton, J. M. Diversity of protein-protein interactions. EMBO Journal 22, 3486–3492 (2003).

23. Perutz, M. F. & Raidt, H. Stereochemical basis of heat stability in bacterial ferredoxins and in haemoglobin A2. Nature 255, 256–259 (1975).

24. Perutz, M. F. Electrostatic effects in proteins. Science 201, 1187–1191 (1978).

25. Zhou, H. X. & Pang, X. Electrostatic Interactions in Protein Structure, Folding, Binding, and Condensation. Chemical Reviews 118, 1691–1741 (2018).

26. Kamieniarz, K. & Schneider, R. Tools to Tackle Protein Acetylation. Chem. Biol. 16, 1027–1029 (2009).

27. Figaro, S. et al. *Bacillus subtilis* mutants with knockouts of the genes encoding ribonucleases RNase Y and RNase J1 are viable, with major defects in cell morphology, sporulation, and competence. J. Bacteriol. 195, 2340–8 (2013).

28. Ducret, A., Quardokus, E. M. & Brun, Y. V. MicrobeJ, a tool for high throughput bacterial cell detection and quantitative analysis. Nat. Microbiol. 1, 16077 (2016).

29. El Mortaji, L., et al. A peptide of a type I toxin-antitoxin system induces *Helicobacter pylori* morphological transformation from spiral shape to coccoids. Proc. Natl. Acad. Sci. U. S. A. 117, 31398–31409 (2020).

30. Christensen, D. G. et al. Post-translational Protein Acetylation: An Elegant Mechanism for Bacteria to Dynamically Regulate Metabolic Functions. Front. Microbiol. 10, 1604 (2019).

31. Christensen, D. G. et al. Identification of Novel Protein Lysine Acetyltransferases in *Escherichia coli*. MBio 9, e01905–18 (2018).

32. Christensen, D. G. et al. Mechanisms, Detection, and Relevance of Protein Acetylation in Prokaryotes. MBio 10, 02708–18 (2019).

33. Hausmann, S. et al. Both exo- and endo-nucleolytic activities of RNase J1 from *Staphylococcus aureus* are manganese dependent and active on triphosphorylated 5′-ends. RNA Biol. 14, 1431–1443 (2017).

34. Tejada-Arranz, A. et al. RNase R is associated in a functional complex with the RhpA DEAD-box RNA helicase in *Helicobacter pylori*. Nucleic Acids Res. 49, 5249–5264 (2021).

35. El Ghachi, M., et al. Characterization of the elongasome core PBP2: MreC complex of *Helicobacter pylori*. Mol. Microbiol. 82, 68–86 (2011).

36. Isono, K. & Isono, S. Ribosomal protein modification in *Escherichia coli*. II. Studies of a mutant lacking the N-terminal acetylation of protein S18. Mol. Gen. Genet. 177, 645–51 (1980).

37. Pathak, D., Bhat, A. H., Sapehia, V., Rai, J. & Rao, A. Biochemical evidence for relaxed substrate specificity of Nα-acetyltransferase (Rv3420c/rimI) of *Mycobacterium tuberculosis*. Sci. Rep. 6, 28892 (2016).

38. Sievers, F. et al. Fast, scalable generation of high-quality protein multiple sequence alignments using Clustal Omega. Mol. Syst. Biol. 7, 539 (2011).

39. Farnbacher, M. et al. Sequencing, annotation, and comparative genome analysis of the gerbil-adapted *Helicobacter pylori* strain B8. BMC Genomics 11, 335 (2010).

40. McClain, M. S., Shaffer, C. L., Israel, D. A., Peek, R. M. & Cover, T. L. Genome sequence analysis of *Helicobacter pylori* strains associated with gastric ulceration and gastric cancer. BMC Genomics 10, 3 (2009).

41. Gibson, D. G. et al. Enzymatic assembly of DNA molecules up to several hundred kilobases. Nat. Methods 6, 343–345 (2009).

42. Karimova, G., Pidoux, J., Ullmann, A. & Ladant, D. A bacterial two-hybrid system based on a reconstituted signal transduction pathway. Proc. Natl. Acad. Sci. U. S. A. 95, 5752–6 (1998).

43. Winn, M. D. et al. Overview of the CCP4 suite and current developments. Acta Crystallogr. D. Biol. Crystallogr. 67, 235–42 (2011).

44. McCoy, A. J. et al. Phaser crystallographic software. J. Appl. Crystallogr. 40, 658–674 (2007).

45. Emsley, P., Lohkamp, B., Scott, W. G. & Cowtan, K. Features and development of Coot. Acta Crystallogr. D. Biol. Crystallogr. 66, 486–501 (2010).

46. Afonine, P. V et al. Towards automated crystallographic structure refinement with phenix.refine. Acta Crystallogr. D. Biol. Crystallogr. 68, 352–67 (2012).

47. Kelley, L. A., Mezulis, S., Yates, C. M., Wass, M. N. & Sternberg, M. J. E. The Phyre2 web portal for protein modeling, prediction and analysis. Nat. Protoc. 10, 845–858 (2015).

48. Krissinel, E. & Henrick, K. Inference of Macromolecular Assemblies from Crystalline State. J. Mol. Biol. 372, 774–797 (2007).

49. Miller, J. H. Experiments in molecular genetics. Cold Spring Harbor Laboratory (1972).

50. Schuck, P. Size-distribution analysis of macromolecules by sedimentation velocity ultracentrifugation and Lamm equation modeling. Biophys. J. 78, 1606–1619 (2000).

51. Laue, T., Shah, B., Ridgeway, T. & Pelletier, S. Computer-aided interpretation of analytical sedimentation data for proteins. in Analytical Ultracentrifugation in Biochemistry and Polymer Science (eds. Harding, S. E., Rowe, A. & Horton, J.) 90–125 Cambridge: Royal Society of Chemistry, (1992).

52. Ortega, A., Amorós, D. & García de la Torre, J. Prediction of hydrodynamic and other solution properties of rigid proteins from atomic- and residue-level models. Biophys. J. 101, 892–8 (2011).

53. Micsonai, A. et al. Accurate secondary structure prediction and fold recognition for circular dichroism spectroscopy. Proc. Natl. Acad. Sci. U. S. A. 112, E3095–103 (2015).

54. Micsonai, A. et al. BeStSel: a web server for accurate protein secondary structure prediction and fold recognition from the circular dichroism spectra. Nucleic Acids Res. 46, W315–W322 (2018).

55. Rappsilber, J., Mann, M. & Ishihama, Y. Protocol for micro-purification, enrichment, pre-fractionation and storage of peptides for proteomics using StageTips. Nat. Protoc. 2, 1896–906 (2007).

56. Tyanova, S., Temu, T. & Cox, J. The MaxQuant computational platform for mass spectrometry-based shotgun proteomics. Nat. Protoc. 11, 2301– 2319 (2016).

57. Cox, J. et al. Andromeda: a peptide search engine integrated into the MaxQuant environment. J. Proteome Res. 10, 1794–805 (2011).

58. MacLean, B. et al. Skyline: an open source document editor for creating and analyzing targeted proteomics experiments. Bioinformatics 26, 966–8 (2010).

59. Perez-Riverol, Y. et al. The PRIDE database and related tools and resources in 2019: improving support for quantification data. Nucleic Acids Res. 47, D442–D450 (2019).

