## Supplemental figures and tables for "Acetylation regulates the oligomerization state and activity of RNase J, the major ribonuclease of *Helicobacter pylori*"

**Supplementary figures**

**Supp Figure S1.** Coomassie blue-stained gels presenting the purification of wild type RNase J and RhpA proteins and of the different RNase J variants used in this study.

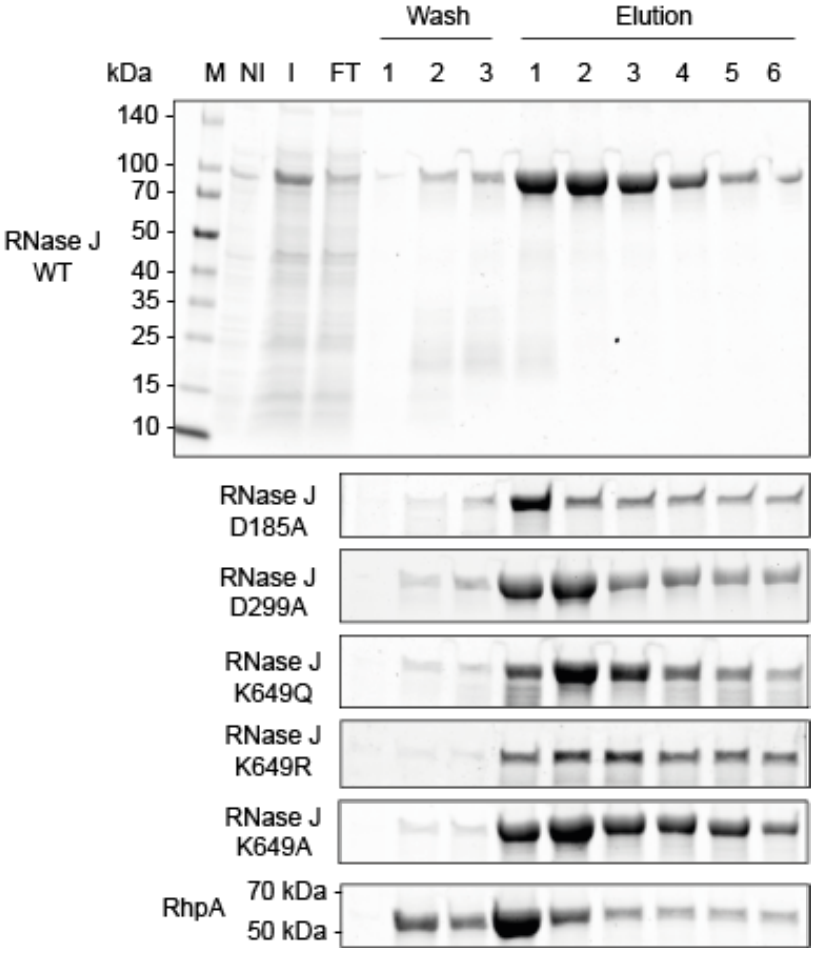

**Supp Figure S2.** Linear map of RNase J showing its domains and the positions of the different acetylated lysine residues detected in this study. NTD: N-terminal domain, CTD: C-terminal domain.

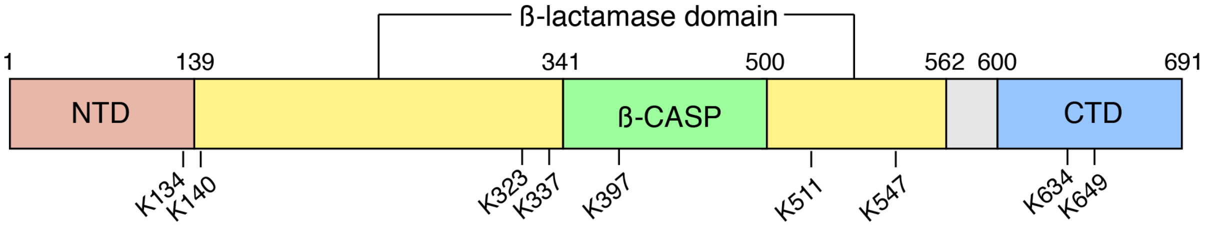

**Supp Figure S3.** Circular dichroism spectra of RNase J WT and variant K649A.

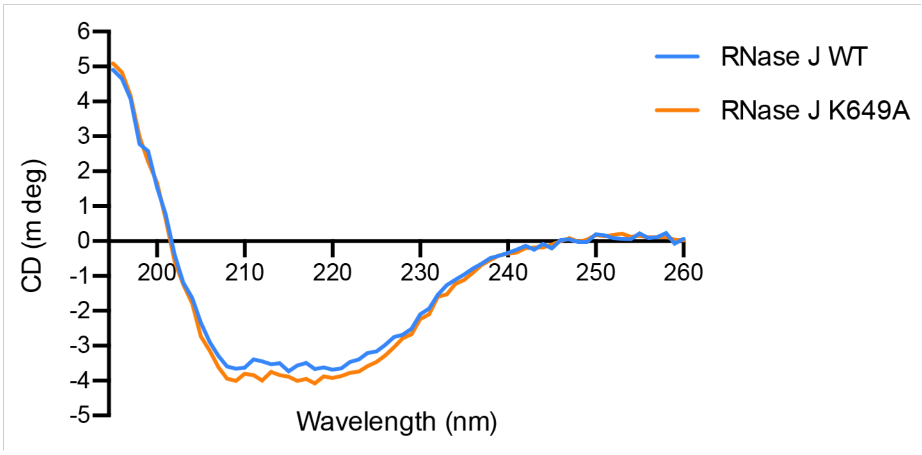

**Supp Figure S4.** (A) Secondary structure of the 45-nt RNA substrate labeled with 6-carboxyfluorescein (6-FAM) at its 3'-end used to assess the activity of the RNase J-RhpA complex. (B) Activity of the RNase J using RNase J WT and the different mutants (D299A, K649A, K649R, K649Q and D185A) over the course of 75 min. (C) Activity of the RNase J-RhpA complex using RNase J WT and the K the different mutants (D299A, K649A, K649R, K649Q and D185A) over the course of 30 min.

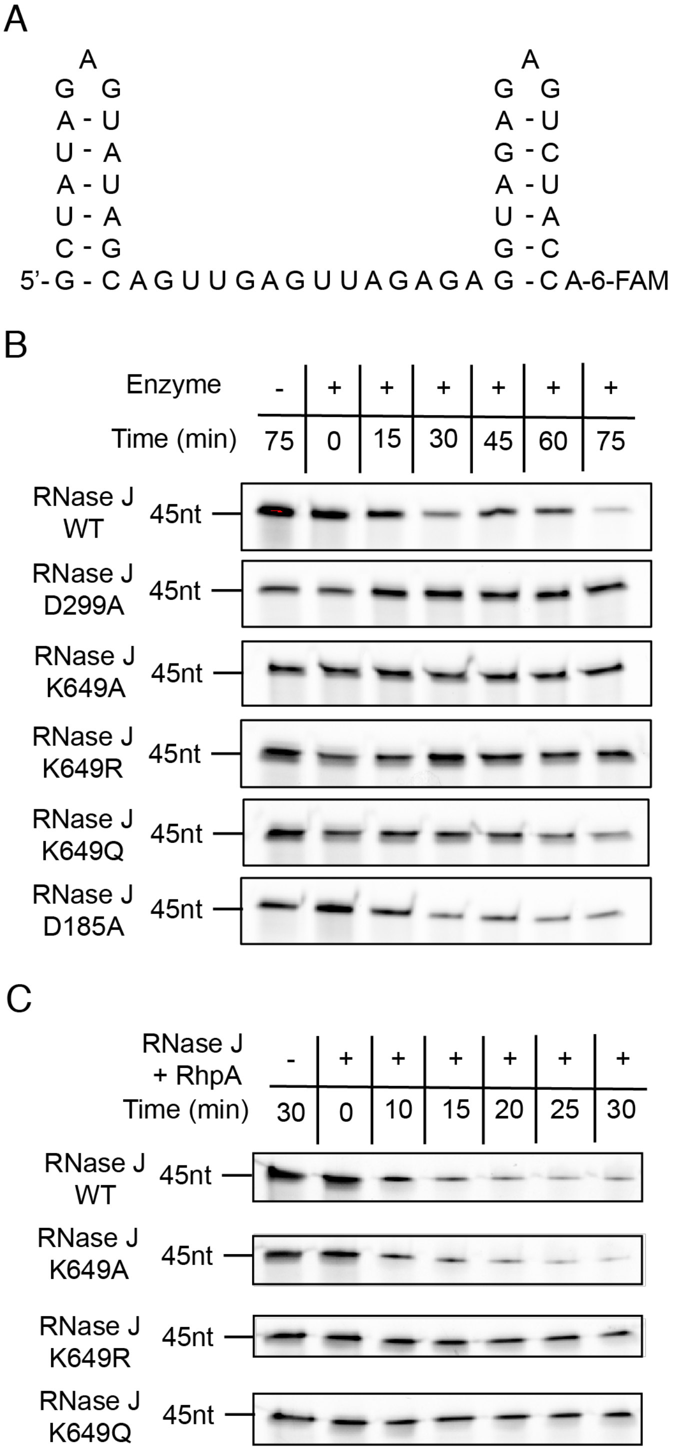

**Supp Figure S5:** Alignment of the RNase J proteins from *H. pylori*, *C. jejuni*, *S. pyogenes*, *D. radiodurans* and the RNases J1 from *B. subtilis* and *S. aureus*. Highlighted in yellow are the acetylated lysine residues from *H. pylori* RNase J and their possible counterparts in the other RNase J sequences.

```
CLUSTAL O(1.2.4) multiple sequence alignment


DraRNase      ------------------------------------------------------------   0
SpyRNase      ------------------------------------------------------------   0
BsuRNase      ------------------------------------------------------------   0
SauRNase      ------------------------------------------------------------   0
HpRNase       MTDNNHYENNESNENSSENSKVDEARAGAFERFTNRKKRFRENAQKNGESSHHEAPSHHK   60
CpyRNase      ---------------------MNEENKTPVAERTNRKHRYKHH-----------------   22

DraRNase      ------------------------------------------------------------   0
SpyRNase      ------------------------------------------------------------   0
BsuRNase      ------------------------------------------------------------   0
SauRNase      ------------------------------------------------------------   0
HpRNase       KEHRPNKKPNNHHKQKHAKTRNYAKEELDSN----KV------EGVTEILHVNERGTLGF   110
CpyRNase      ---RD-----NLKKQANANTQNVQANEVATDIAESEVKKPKKRKHKNGNNTVKISGNEGW   74

DraRNase      --------------MTRPEQPRPESADLPAPTLEVIPLGGMGEIGKNITVFRYGDEIVVV   46
SpyRNase      ----------------------MTNISLKPNEVGVFAIGGLGEIGKNTYGIEYQDEIIIV   38
BsuRNase      ------------------------MKFVKNDQTAVFALGGLGEIGKNTYAVQFQDEIVLI   36
SauRNase      ------------------------MKQLHPNEVGVYALGGLGEIGKNTYAVEYKDEIVII   36
HpRNase       HKELKKGVETNNKIQVEHLNPHYKMNLNSKASVKITPLGGLGEIGGNMMVIETPKSAIVI   170
CpyRNase      QKDMQASIEANRASHELRLNPLKYL-NSSEHKIKITPLGGLGEIGGNMTVFETDNDAIIV   133
                                                ::  :**:**** *   ..  .. :::

DraRNase      DGGLAFPKAHQMGIDLIVPRIDYLLEHQDKIKGWILTHGHEDHIGGLPYIFARLPRVPVY   106
SpyRNase      DAGIKFPEDDLLGIDYVIPDYSYIVDNLDRVKALVITHGHEDHIGGIPFLLKQA-NIPIY   97
BsuRNase      DAGIKFPEDELLGIDYVIPDYTYLVKNEDKIKGLFITHGHEDHIGGIPYLLRQV-NIPVY   95
SauRNase      DAGIKFPDDNLLGIDYVIPDYTYLVQNQDKIVGLFITHGHEDHIGGVPFLLKQL-NIPIY   95
HpRNase       DAGMSFPKEGLFGVDILIPDFSYLHQIKDKIAGIIITHAHEDHIGATPYLFKEL-QFPLY   229
CpyRNase      DIGMSFPSESMHGVDILIPDFDYIRKIKQKVRGIIITHAHEDHIGAVPYFFKEF-QFPIY   192
              * *: **.    *:* ::*   *: .  ::: . .:**.******. *::: .  ..*:*

DraRNase      GLPLTLALVREKLSEFGLQDVDLR-EVTYGDEVRFGQSFVAEFFCMTHSIPDNAGYILKT   165
SpyRNase      AGPLALALIRGKLEEHGLWREATVYEI-NHNTELTFKNMSVTFFKTTHSIPEPVGIVIHT   156
BsuRNase      GGKLAIGLLRNKLEEHGLLRQTKLNII-GEDDIVKFRKTAVSFFRTTHSIPDSYGIVVKT   154
SauRNase      GGPLALGLIRNKLEEHHLLRTAKLNEI-NEDSVIKSKHFTISFYLTTHSIPETYGVIVDT   154
HpRNase       GTPLSLGLIGSKFDEHGLKKYRSYFKIVEKRCPISVGEFIIEWIHITHSIIDSSALAIQT   289
CpyRNase      ATPLPLGMISNKFEEHGLKAHRSYFRPIEKRKLYEIGDFEIELIHITHSIIDASALVIIT   252
              .  * :.::  *:.*. *                            **** :  .  : *

DraRNase      PVGDVLHTGDFKIDPDVGTGAGIVSDLERVEQAGKDGVLLLISDSTNAERPGHTPSEAEI   225
SpyRNase      PQGKIICTGDFKFDFTPV---GDPADLQRMAALGEEGVLCLLSDSTNAEIPTFTNSEKVV   213
BsuRNase      PPGNIVHTGDFKFDFTPV---GEPANLTKMAEIGKEGVLCLLSDSTNSENPEFTMSERRV   211
SauRNase      PEGKVVHTGDFKFDFTPV---GKPANIAKMAQLGEEGVLCLLSDSTNSLVPDFTLSEREV   211
HpRNase       KAGTIIHTGDFKIDHTPV--DNLPTDLYRLAHYGEKGVMLLLSDSTNSHKSGTTPSESTI   347
CpyRNase      KAGTILHTGDFKIDHTPI--DGYPTDLNRLAYYGERGVLCMLSDSTNSYKEGITKSESSV   310
                * :: *****:*       .  ::: ::   *: **: ::*****:     * **  :

DraRNase      ARNLEEIIKGCRGRVFLTTFASQVYRIQNILDLAHRQGRRVVMEGRSMIKYAQAAQATGH   285
SpyRNase      GQSILKIIEGIHGRIIFASFASNIYRLQQAAEAAVKTGRKIAVFGRSMEKAIVNGIELGY   273
BsuRNase      GESIHDIFRKVDGRIIFATFASNIHRLQQVIEAAVQNGRKVAVFGRSMESAIEIGQTLGY   271
SauRNase      GQNVDKIFRNCKGRIIFATFASNIYRVQQAVEAAIKNNRKIVTFGRSMENNIKIGMELGY   271
HpRNase       APAFDTLFKEAQGRVIMSTFSSNIHRVYQAIQYGIKYNRKIAVIGRSMEKNLDIARELGY   407
CpyRNase      GKTFDAIFATSKGRVIMSTFSSNIHRVYQAIERGVKHGRKVCVIGRSMERNLWTAIELGY   370
              .  .  ::    **:::::*:*:::*: :  : . : .*::   ****      .   *:

DraRNase      MN-PPEPFLTSEEVGELQDQQVLFVCTGSQGQPMAVLGRLAFGTHAKIALRRGDTVILSS   344
SpyRNase      IKVPKGTFIEPSELKNLHASEVLIMCTGSQGESMAALARIANGTHRQVTLQPGDTVIFSS   333
BsuRNase      INCPKNTFIEHNEINRMPANKVTILCTGSQGEPMAALSRIANGTHRQISINPGDTVVFSS   331
SauRNase      IKAPPETFIEPNKINTVPKHELLILCTGSQGEPMAALSRIANGTHKQIKIIPEDTVVFSS   331
HpRNase       IHLPYQSFIEANEVAKYPDNEVLIVTTGSQGETMSALYRMATDEHRHISIKPNDLVIISA   467
CpyRNase      VNLDKKIFIDANEVSKYPDNEVLIVTTGSQGETMSALYRMATDEHKYIKIKPTDQIIISS   430
              ::     *:  .::      :: :: *****: *:.* *:* . *  : :   * :::*:
```

```
DraRNase        NPIPGNEDAVNLIVNRLYEIGVDVVYPPTYRVHASGHASQEELATILNLTRPKFFLPWHG  404
SpyRNase        SPIPGNTTSVNKLINTIQEAGVDVIHGKVNNIHTSGHGGQQEQKLMLSLIKPKYFMPVHG  393
BsuRNase        SPIPGNTISVSRTINQLYRAGAEVIHGPLNDIHTSGHGGQEEQKLMLRLIKPKFFMPIHG  391
SauRNase        SPIPGNTKSINRTINSLYKAGADVIHSKISNIHTSGHGSQGDQQLMLRLIKPKYFLPIHG  391
HpRNase         KAIPGNEASVSAVLNFLIKKEAKVAYQEFDNIHVSGHAAQEEQKLMLRLIKPKFFLPVHG  527
CpyRNase        KAIPGNETSVSTVLNYLLKSGASVAHQDFSEIHVSGHAAQEEQKLMLRLVKPKFFLPVHG  490
                . ****  ::.  :* : .  ..* :     :*.***..* :   :* * :**:*:* **


DraRNase        EPRHQINHAKLAQTLPRPPKRTLIAKNGDIVNLGPDEFRVSGTVAAGAVYVDGLGVGDVN  464
SpyRNase        EYRMQKIHAGLAMDIGIPKENIFIMENGDVLALTSDSARIAGHFNAQDIYVDGNGIGDIG  453
BsuRNase        EYRMQKMHVKLATDCGIPEENCFIMDNGEVLALKGDEASVAGKIPSGSVYIDGSGIGDIG  451
SauRNase        EYRMLKAHGETGVECGVEEDNVFIFDIGDVLALTHDSARKAGRIPSGNVLVDGSGIGDIG  451
HpRNase         EYNHVARHKQTAISCGVPEKNIYLMEDGDQVEVGPAFIKKVGTIKSGKSYVDNQSNLSID  587
CpyRNase        EYNHIVRHKETAIACGVDERNTYLMSDGDQIEVCQKYIKRLKTVKTGKVFIDNQINKQIS  550
                * .    *   .        .  : . *: : :          . :    :*.    .:.

DraRNase        DDVLLDRVNLSQEGLLILTAVLHPTPH-----VEVVARGFAR--PNRDLELQIRRVALEA  517
SpyRNase        AAVLRDRRDLSEDGVVLAVATVDFNTQMILAGPDILSRGFIYMRESGDLIRESQRVLFNA  513
BsuRNase        NIVLRDRRILSEEGLVIVVVSIDMDDFKISAGPDLISRGFVYMRESGDLINDAQELISNH  511
SauRNase        NVVIRDRKLLSEEGLVIVVVSIDFNTNKLLSGPDIISRGFVYMRESGQLIYDAQRKIKTD  511
HpRNase         TSIVQQREEVASAGVFAATIFVNKNKQALLESSQFSSLGLVGFKDEKHLIKEIQGGLEML  647
CpyRNase        DDVVIDRQKLAEAGVVTIISQIDKNAKTLIQN-RVISYGLVSERQSKNLSKEMEEVLLQF  609
                  :: :*  ::. *:.     :.           . : *:     . .*  : .

DraRNase        VEQGLR--EKKRLEDVRDDMYGAVRRFTRKATGRNPVLIPMIVD----------        559
SpyRNase        -IRIALKNKDASIQSVNGAIVNALRPFLYEKTEREPIIIPMVLTPDKH------        560
BsuRNase        -LQKVMERKTTQWSEIKNEITDTLAPFLYEKTKRRPMILPIIMEV---------        555
SauRNase        VISKLNQNKDIQWHQIKSSIIETLQPYLFEKTARKPMILPVIMKVNEQKESNNK        565
HpRNase         L-KSSNAEILNNPKKLEDHTRNFIRKALFKKFRKYPAIICHAHSF---------        691
CpyRNase        L-SNVKDELLHDQRALENQIRQVIRKHIFRKIKKYPTIVPVVYLM---------        653
                               :..     :     .   : * ::
```

**Supplementary Tables**

**Supp Table S1**: RNase J crystallographic data collection and refinement statistics.

| | **RNase J core** |
|---|---|
| **Wavelength** | 0.97778 |
| **Resolution range** | 56.03 - 2.75 (2.848 - 2.75) |
| **Space group** | I 41 2 2 |
| **Unit cell** | 158.49 158.49 214.26 90 90 90 |
| **Total reflections** | 128437 |
| **Unique reflections** | 35426 (3469) |
| **Multiplicity** | 3.6 (3.6) |
| **Completeness (%)** | 99.17 (99.06) |
| **Mean I/sigma(I)** | 5.9 (1.3) |
| **Wilson B-factor** | 75.00 |
| **R-merge** | 0.137 (0.80) |
| **R-meas** | 0.160 (0.94) |
| **R-pim** | 0.081 (0.486) |
| **CC1/2** | 0.979 (0.453) |
| **Reflections used in refinement** | 35419 (3467) |
| **Reflections used for R-free** | 1805 (165) |
| **R-work** | 0.2009 (0.3197) |
| **R-free** | 0.2318 (0.3620) |
| **Number of non-hydrogen atoms** | 3559 |
| **macromolecules** | 3530 |
| **solvent** | 29 |

| | |
|---|---|
| **Protein residues** | 450 |
| **RMS(bonds)** | 0.009 |
| **RMS(angles)** | 1.05 |
| **Ramachandran favored (%)** | 92.86 |
| **Ramachandran allowed (%)** | 6.47 |
| **Ramachandran outliers (%)** | 0.67 |
| **Rotamer outliers (%)** | 2.07 |
| **Clashscore** | 6.06 |
| **Average B-factor** | 72.57 |
| **macromolecules** | 72.62 |
| **solvent** | 67.25 |

Statistics for the highest-resolution shell are shown in parentheses.

**Supp Table S2.** Mass spectrometry data of the raw intensity of the acetylated RNase J peptides, the peptides used for normalization across conditions and the normalized data.

**Supp Table S3.** Estimation of the proportions of the secondary structure elements of wild type RNase J and variant K649A from their circular dichroism spectra.

| Secondary structure | RNase J WT (%) | RNase J K649A (%) |
|---|---|---|
| Alpha-helix | 4.4 | 5.2 |
| Parallel beta-sheet | 0 | 0 |
| Antiparallel beta-sheet | 37.5 | 36.9 |
| Turn | 14.4 | 14.5 |
| Others | 43.5 | 43.4 |

**Supp Table S4.** Strains and plasmids used in this study.

| **Strains** | **Relevant characteristics** | **Reference** |
|---|---|---|
| *Escherichia coli* | | |
| XL1-Blue | *recA1 endA1 gyrA96 thi-1 hsdR17 supE44 relA1 lac* [F *proAB lacIqZΔM15 Tn10* (Tetr)] | Commercially available strain (Agilent Technologies) |
| Bli5 | *F- mcrA* Δ( *mrr-hsd* RMS-*mcr*BC) Φ80*lac*ZΔM15 Δ*lac*X74 *rec*A1 *ara*D139 Δ(*ara leu*)7697 *gal*U *gal*K *rps*L (StrR) *end*A1 *nup*G, $Cm^R$ | (Munier et al., 1991) |
| BTH101 | Strain for two hybrid assays | (Karimova, Pidoux, Ullmann, & Ladant, 1998) |
| *Helicobacter pylori* | | |
| B128 | Sequenced parental strain | (Farnbacher et al., 2010; McClain, Shaffer, Israel, Peek, & Cover, 2009) |
| B128 RNase J-FLAG | $Apra^R$, fusion protein RNase J-FLAG | (Tejada-Arranz et al., 2020) |
| B128 RNase J-FLAG Δ*rimI::apra* | $Cm^R$, $Apra^R$ | This work |
| B128 RNase J-FLAG Δ*hpb8_1270::apra* | $Cm^R$, $Apra^R$ | This work |
| B128 RNase J-FLAG Δ*pta-ackA::apra* | $Cm^R$, $Apra^R$ | This work |
| B128 pILL2157-RNase J | $Cm^R$ | This work |
| B128 pILL2157-RNase J K134A | $Cm^R$ | This work |
| B128 pILL2157-RNase J K323A | $Cm^R$ | This work |
| B128 pILL2157-RNase J K337A | $Cm^R$ | This work |
| B128 pILL2157-RNase J K397A | $Cm^R$ | This work |

| | | |
|---|---|---|
| B128 pILL2157-RNase J K511A | $Cm^R$ | This work |
| B128 pILL2157-RNase J K547A | $Cm^R$ | This work |
| B128 pILL2157-RNase J K634A | $Cm^R$ | This work |
| B128 pILL2157-RNase J K649A | $Cm^R$ | This work |
| B128 pILL2157-RNase J D185A | $Cm^R$ | This work |
| B128 pILL2157-RNase J Δ*rnj::km* | $Cm^R$, $Km^R$ | This work |
| B128 pILL2157-RNase J K134A Δ*rnj::km* | $Cm^R$, $Km^R$ | This work |
| B128 pILL2157-RNase J K323A Δ*rnj::km* | $Cm^R$, $Km^R$ | This work |
| B128 pILL2157-RNase J K337A Δ*rnj::km* | $Cm^R$, $Km^R$ | This work |
| B128 pILL2157-RNase J K397A Δ*rnj::km* | $Cm^R$, $Km^R$ | This work |
| B128 pILL2157-RNase J K511A Δ*rnj::km* | $Cm^R$, $Km^R$ | This work |
| B128 pILL2157-RNase J K547A Δ*rnj::km* | $Cm^R$, $Km^R$ | This work |
| B128 pILL2157-RNase J K634A Δ*rnj::km* | $Cm^R$, $Km^R$ | This work |
| B128 pILL2157-RNase J K649A Δ*rnj::km* | $Cm^R$, $Km^R$ | This work |
| B128 pILL2157-RNase J D185A Δ*rnj::km* | $Cm^R$, $Km^R$ | This work |
| B128 pILL2157-RNase J Δ*rnj::km* Δ*rhpA::apra* | $Cm^R$, $Km^R$, $Apra^R$ | (El Mortaji et al., 2018) |
| **Plasmid** | **Resistance** | **Reference** |

| | | |
|---|---|---|
| pET28(a+) | $Km^R$ | Commercial plasmid (Novagen) |
| pET28-RNase J | $Km^R$ | This work |
| pET28-RNase J K649A | $Km^R$ | This work |
| pET28-RNase J K649R | $Km^R$ | This work |
| pET28-RNase J K649Q | $Km^R$ | This work |
| pET28-RNase J D185A | $Km^R$ | This work |
| pET28-RNase J D299A | $Km^R$ | This work |
| pET28-RhpA | $Km^R$ | This work |
| pILL2157 | $Cm^R$ | (Boneca et al., 2008) |
| pILL2157-RNase J | $Cm^R$ | This work |
| pILL2157-RNase J K134A | $Cm^R$ | This work |
| pILL2157-RNase J K323A | $Cm^R$ | This work |
| pILL2157-RNase J K337A | $Cm^R$ | This work |
| pILL2157-RNase J K397A | $Cm^R$ | This work |
| pILL2157-RNase J K511A | $Cm^R$ | This work |
| pILL2157-RNase J K547A | $Cm^R$ | This work |
| pILL2157-RNase J K649A | $Cm^R$ | This work |
| pILL2157-RNase J D185A | $Cm^R$ | This work |
| pUT18 | $Amp^R$ | (Karimova, Pidoux, Ullmann, & Ladant, 1998) |
| pKNT25 | $Km^R$ | (Karimova, Pidoux, Ullmann, & Ladant, 1998) |
| pUT18-RNase J | $Amp^R$ | This work |
| pKNT25-RNase J | $Km^R$ | This work |
| pKNT25-RNase J ΔCt | $Km^R$ | This work |
| pKNT25-RNase J K611A | $Km^R$ | This work |
| pKNT25-RNase J E618A | $Km^R$ | This work |
| pKNT25-RNase J E634A | $Km^R$ | This work |
| pKNT25-RNase J E654A | $Km^R$ | This work |
| pKNT25-RNase J E660A | $Km^R$ | This work |

| | | |
|---|---|---|
| pKNT25-RNase J E663A | $Km^R$ | This work |
| pKNT25-RNase J E595A | $Km^R$ | This work |
| pKNT25-RNase J E596A | $Km^R$ | This work |

**Supp Table S5:** oligonucleotides used in this study.

| Name | Sequence | Use |
|---|---|---|
| Chromosomal deletions | | |
| oATA267 | TTTGATTTCATTCTT ACAGG | fw oligo to amplify the upstream region to hpb8_615 |
| oATA268 | GTATTGCACGACATT GCACTCCGATTAACT TTTTAAAACCC | rv oligo to amplify the upstream region to hpb8_615, with a region complementary to the beginning of the apramycin cassette |
| oATA269 | GCTGATACCTGGAG GGAATATTGGCGCA TCCATTTGACACACG | fw oligo to amplify the downstream region to hpb8_615, with a region complementary to the end of the non-polar apramycin cassette |
| oATA270 | GTTTTCTATAGTTAA TTGACC | rv oligo to amplify the downstream region to hpb8_615 |
| oATA271 | CGTTTGCCAGAATTA GAGCC | fw oligo to amplify the upstream region to hpb8_1270 |
| oATA272 | GTATTGCACGACATT GCACTCCAACAACT CTCTTTTATTTCTTTA AC | rv oligo to amplify the upstream region to hpb8_1270 with a region complementary to the beginning of the apramycin cassette |
| oATA273 | GCTGATACCTGGAG GGAATAATGATTTTT GGGGATTTTAAATAT C | fw oligo to amplify the downstream region to hpb8_1270 with a region complementary to the end of the apramycin cassette |
| oATA274 | AATTCAAAAAACAAT TCCGG | rv oligo to amplify the downstream region to hpb8_1270 |
| oLEM114 | GGAGTGCAATGTCG TGCAATAC | Fw oligo to amplify the apramycin resistance cassette |
| oATA243 | TATTCCCTCCAGGTA TCAGCCAATCGACT GGCGAG | rv oligo to amplify the apramycin cassette with an RBS for non-polar deletions |
| oATA286 | TAGGGTTGGGGTTT TTTCAAGC | fw oligo to amplify 500 bp upstream from the beginning of pta |
| oATA287 | GTATTGCACGACATT GCACTCCGCCAATC TCCCTAAAATTATC | rv oligo to amplify 500 bp upstream from the beginning of pta, with a region complementary to the beginning of the apramycin cassette |
| oATA292 | GCTGATACCTGGAG GGAATAAGGTTGTTT AATGAAAATGGTTC | fw oligo to amplify 500 bp downstream from the end of ackA, with a region complementary to the end of the apramycin cassete |
| oATA293 | AAATTTTCGTATCGT AGCGC | rv oligo to amplify 500 bp downstream from the end of ackA |
| oLEM009 | GGTACCCGGGTGAC TAAC | Fw oligo to amplify the kanamycin resistance cassette |
| oLEM010 | CATTATTCCCTCCAG GTAC | Rv oligo to amplify the kanamycin resistance cassette for non-polar deletions |

| pILL2157 constructs | | |
|---|---|---|
| oATA470 | TTTGTTTGGAAGGAAAAGCAATGACTGATAACAACCATTATGAAAACAATGAAAGC | fw oligo to clone RNase J into pILL2157, digested with *Nde*I, by isoT assembly |
| oATA471 | ATTCGAGCTCGGTACCCGGGTCAAAAAGAATGGGCATGACAAATGATAGCC | rv oligo to clone RNase J into pILL2157, digested with *Bam*HI, by isoT assembly |
| oATA400 | ATGAATCTAAACTCTAAAGC | fw oligo to introduce mutations in position 134 in RNase J |
| oATA403 | GCTTTAGAGTTTAGATTCATCGCATAATGCGGGTTTAGATGC | rv oligo to introduce mutation K134A, aag402gcg, in RNase J |
| oATA517 | GCGAGCGTTAAAATCACGCC | fw oligo to introduce mutations in residue 140 of RNase J |
| oATA520 | GGCGTGATTTTAACGCTCGCTGCAGAGTTTAGATTCATCTTATAATG | rv oligo to introduce mutation K140A, aaa420gca, in RNase J |
| oATA404 | GGGGTGATGCTTCTTTTAAGC | fw oligo to introduce mutations in position 323 in RNase J |
| oATA407 | GCTTAAAAGAAGCATCACCCCCGCTTCGCCATAGTGCGCTAAACG | rv oligo to introduce mutation K323A, aag969gcg, in RNase J |
| oATA408 | TCCGGGACCACGCCGAGTGAAAGC | fw oligo to introduce mutations in position 337 in RNase J |
| oATA411 | GCTTTCACTCGGCGTGGTCCCGGATGCATGGGAGTTAGTGGAATCG | rv oligo to introduce mutation K337A, aaa1011gca, in RNase J |
| oATA412 | AACCTAGACATCGCCAGAGAATTAGG | fw oligo to introduce mutations in position 397 in RNase J |
| oATA415 | CCTAATTCTCTGGCGATGTCTAGGTTTGCTTCCATAGAACGCCCGATCAC | rv oligo to introduce mutation K397A, aaa1191gca, in RNase J |
| oATA416 | CTCATGTTAAGACTCATTAAGC | fw oligo to introduce mutations in position 511 in RNase J |
| oATA419 | GCTTAATGAGTCTTAACATGAGCGCTTGCTCTTCTTGAGCGG | rv oligo to introduce mutation K511A, aag1533gcg, in RNase J |
| oATA521 | CATTTGATCAAAGAAATTCAAGG | fw oligo to introduce mutations in residue 634 of RNase J |
| oATA524 | CCTTGAATTTCTTTGATCAAATGCGCTTCA | rv oligo to introduce mutation K634A, aag1902gcg, in RNase J |

| | | |
|---|---|---|
| | TCTTTAAAGCCCACAAGC | |
| oATA297 | TGCCAATAACATTTCTAATCCCC | rv oligo to amplify rnj 1-648 with extra nt to introduce mutation K649A, aaa1947gca |
| oATA298 | GGGGATTAGAAATGTTATTGGCATCCAGCAACGCCGAAATTTTG | fw oligo to amplify rnj 650-691 with an overlapping region to oATA297 to introduce mutation K649A, aaa1947gca |
| oATA380 | AGCCACGCCAAATAGCCCCTC | rv oligo to amplify rnj 1-184 with extra nt to introduce mutation D185A, gat555gct |
| oATA381 | GAGGGGCTATTTGGCGTGGCTATTTTAATCCCGGATTTTTCC | fw oligo to amplify rnj 186-691 with an overlapping region to oATA380 to introduce mutation D185A, gat555gct |
| pET28 constructs | | |
| oATA396 | ATATCATATGGAGAATCTTTATTTTCAAGGAATGACTGATAACAACCATTATG | fw oligo to amplify rnj to introduce it in pET28a+ with NdeI restriction site and including a TEV restriction site |
| oATA397 | AGAGCTCGAGTCAAAAAGAATGGGCATGACAAATG | rv oligo to amplify rnj to introduce it in pET28a+ with XhoI restriction site |
| oATA420 | TCCAGCAACGCCGAAATTTTG | fw oligo to introduce mutations in position 649 in RNase J |
| oATA421 | CAAAATTTCGGCGTTGCTGGATCTCAATAACATTTCTAATCC | rv oligo to introduce mutation K649R, aaa1947aga, in RNase J |
| oATA422 | CAAAATTTCGGCGTTGCTGGATTGCAATAACATTTCTAATCC | rv oligo to introduce mutation K649Q, aaa1947caa, in RNase J |
| oATA451 | atatCATATGGAGAATCTTTATTTTCAAGGAATGGAATTGAATCAACCACCACTCC | fw oligo to amplify rhpA to introduce it in pET28a+ with NdeI restriction site and including a TEV restriction site |
| oATA090 | AGAGCTCGAGTTAACGGCGTTTGGGTTTTTTAGAATGG | rv oligo to amplify rhpA to introduce it in pET28a+ with STOP codon and XhoI restriction site |
| pUT18 constructs | | |
| oATA046 | GAGCTGCAGGATGACTGATAACAACCATTATGAAAACAATGAAAGC | fw oligo for cloning *rnj* in pKNT25, pUT18 and pUT18C in frame with T25/T18 in the PstI restriction site |
| oATA047 | CTCGGTACCCGAAAAGAATGGGCATGACAAATGATAGCC | rv oligo for cloning *rnj* in double-hybrid plasmids in KpnI restriction site without the STOP codon, in frame with T25/T18 |
| pKNT25 constructs | | |
| oATA216 | CTCGGTACCCGGCTAGCGACTTCTTCTCTTTGTTGC | rv oligo for cloning *ΔCt-rnj* in two-hybrid plasmids in KpnI restriction site without the STOP codon, in frame with T25/T18 |

| | | |
|---|---|---|
| oATA295 | TGCATTCACAAAAATCGTAGCCGC | rv oligo to amplify rnj 1-610 with extra nt to introduce mutation K611A, aaa1833gca |
| oATA296 | GCGGCTACGATTTTTGTGAATGCAAACAAGCAAGCGCTTTTAG | fw oligo to amplify rnj 612-691 with an overlapping region to oATA295 to introduce mutation K611A, aaa1833gca |
| oATA299 | cgcttcatctttaaagcccacaagc | rv oligo to amplify rnj 1-633 with extra nt to introduce mutation K634A, aag1902gcg |
| oATA300 | GCTTGTGGGCTTTAAAGATGAAGCGCATTTGATCAAAGAAATTCAAGG | fw oligo to amplify rnj 635-691 with an overlapping region to oATA299 to introduce mutation K634A, aag1902gcg |
| oATA301 | TGCAGGGTTATTCAAAATTTCG | rv oligo to amplify rnj 1-659 with extra nt to introduce mutation K660A, aaa1980gca |
| oATA302 | CGAAATTTTGAATAACCCTGCAAAATTAGAAGATCACACTCG | fw oligo to amplify rnj 661-691 with an overlapping region to oATA301 to introduce mutation K660A, aaa1980gca |
| oATA303 | TGCTAAAAGCGCTTGCTTGTTTTTATTC | rv oligo to amplify rnj 1-617 with extra nt to introduce mutation E618A, gaa1854gca |
| oATA304 | GAATAAAAACAAGCAAGCGCTTTTAGCAAGCTCTCAATTTTCCAGTTTAGG | fw oligo to amplify rnj 619-691 with an overlapping region to oATA303 to introduce mutation E618A, gaa1854gca |
| oATA305 | TGCGGCGTTGCTGGATTTCAATAAC | rv oligo to amplify rnj 1-653 with extra nt to introduce mutation E654A, gaa1962gca |
| oATA306 | GTTATTGAAATCCAGCAACGCCGCAATTTTGAATAACCCTAAAAAATTAG | fw oligo to amplify rnj 655-691 with an overlapping region to oATA305 to introduce mutation E654A, gaa1962gca |
| oATA307 | TGCTAATTTTTTAGGGTTATTC | rv oligo to amplify rnj 1-662 with extra nt to introduce mutation E663A, gaa1989gca |
| oATA308 | GAATAACCCTAAAAAATTAGCAGATCACACTCGTAATTTCATC | fw oligo to amplify rnj 664-691 with an overlapping region to oATA307 to introduce mutation E663A, gaa1989gca |
| oATA311 | TGCTCTTTGTTGCACGATGCTTGTG | rv oligo to amplify rnj 1-594 with extra nt to introduce mutation E595A, gaa1785gca |
| oATA312 | CACAAGCATCGTGCAACAAAGAGCAGAAGTCGCTAGCGCCGGGGTG | fw oligo to amplify rnj 596-691 with an overlapping region to oATA311 to introduce mutation E595A, gaa1785gca |
| oATA313 | TGCTTCTCTTTGTTGCACGATGC | rv oligo to amplify rnj 1-595 with extra nt to introduce mutation E596A, gaa1788gca |
| oATA314 | GCATCGTGCAACAAAGAGAAGCAGTCGCTAGCGCCGGGGTGTTTGC | fw oligo to amplify rnj 597-691 with an overlapping region to oATA313 to introduce mutation E596A, gaa1788gca |
| Ribonuclease activity tests (RNA oligos) | | |

| | | |
|---|---|---|
| oMBL001 | 6-FAM-AAUCGUUAUGACUGAUAACAACCA | RNA substrate labeled in 5’ for exoribonuclease activity tests |
| oMBL002 | AAUCGUUAUGACUGAUAACAACCA-6-FAM | RNA substrate labeled in 3’ for exoribonuclease activity tests |
| oMBL004 | GCUAUAGAGUAUAGCAGUUGAGUUAGAGAGGUAGAGAGUCUACCA-6-FAM | RNA substrate labeled in 3’ for endoribonuclease activity tests |
